# Endosomal egress and intercellular transmission of hepatic ApoE-containing lipoproteins and its exploitation by the hepatitis C virus

**DOI:** 10.1101/2022.12.08.519703

**Authors:** Minh-Tu Pham, Ji-Young Lee, Christian Ritter, Roman Thielemann, Uta Haselmann, Charlotta Funaya, Vibor Laketa, Karl Rohr, Ralf Bartenschlager

**Affiliations:** Department of Infectious Diseases, Molecular Virology, Center for Integrative Infectious Diseases Research, Heidelberg University, Heidelberg, Germany; German Center for Infection Research (DZIF), Partner Site Heidelberg, Heidelberg, Germany; Division Virus-Associated Carcinogenesis, German Cancer Research Center (DKFZ), Heidelberg, Germany; BioQuant Center, IPMB, Biomedical Computer Vision Group, Heidelberg University, Heidelberg, Germany; Electron Microscopy Core Facility (EMCF), Heidelberg University, Heidelberg, Germany; Department of Infectious Diseases, Virology, Center for Integrative Infectious Diseases Research, Heidelberg University, Heidelberg, Germany

## Abstract

Liver-generated plasma Apolipoprotein E (ApoE)-containing lipoproteins (LPs) (ApoE-LPs) play central roles in lipid transport and metabolism. Perturbations of ApoE can result in several metabolic disorders and ApoE genotypes have been associated with multiple diseases. ApoE is synthesized at the endoplasmic reticulum and transported to the Golgi apparatus for LP assembly; however, ApoE-LPs transport from there to the plasma membrane is largely unknown. Here, we established an integrative imaging approach based on a fully functional fluorescently tagged ApoE. We found that ApoE-LPs accumulate in CD63-positive endosomes of hepatocytes. In addition, we observed the co-egress of ApoE-LPs and extracellular vesicles (EVs) along the late endosomal trafficking route. Moreover, complexes of ApoE-LPs and CD63-positive EVs were found to be transmitted from cell to cell. Given the important role of ApoE in viral infections, we studied the hepatitis C virus (HCV) and found that the viral replicase protein NS5A is enriched in ApoE-containing intraluminal vesicles. Interaction between NS5A and ApoE is required for the efficient release of EVs containing viral RNA. These vesicles are transported along the endosomal ApoE egress pathway. Taken together, our data argue for endosomal egress and transmission of hepatic ApoE-LPs, a pathway that is hijacked by HCV. Given the more general role of EV-mediated cell-to-cell communication, these insights provide new starting points for research into the pathophysiology of ApoE-related metabolic and infection-related disorders.

**Author Summary:** The post-Golgi egress pathway of hepatocyte-derived ApoE-containing lipoproteins (ApoE-LPs) is largely unknown. By using integrative imaging analyses, we show that ApoE-LPs are enriched in CD63-positive endosomes suggesting that these endosomes might be a central hub for the storage of ApoE-LPs from which they are released into the circulation. In addition, we provide evidence for the co-egress of ApoE-LPs with extracellular vesicles (EVs) along the late endosomal route and their transfer from cell to cell. This pathway is hijacked by the hepatitis C virus that induces the production of ApoE-associated EVs containing viral RNA. Given the important role of ApoE in multiple metabolic, degenerative and infectious diseases, and the role of EVs in cell-to-cell communication, these results provide important information how perturbations of ApoE might contribute to various pathophysiologies.

## Introduction

Hepatocytes play a central role in lipid metabolism, both by production and clearance of plasma lipoproteins (LPs). Changes in hepatic lipid metabolism may contribute to chronic liver disease, such as nonalcoholic fatty liver disease (1). Moreover, infections with hepatotropic viruses, most notably the hepatitis C virus (HCV), perturb hepatic lipid homeostasis, leading to hepatosteatosis, which is due to virus-induced increased intracellular lipid accumulation and impaired lipid release from infected cells (2). These alterations promote viral replication that requires both intracellular lipids to build up a membranous replication factory (3) and to assemble particular virions, designated lipoviroparticles, because of the lipid profile resembling the one of LPs (4) and the association with apolipoproteins, especially apolipoprotein E (ApoE) (5).

LPs such as very-low-density lipoprotein (VLDL) and high-density lipoprotein (HDL) are water-soluble assemblies of macromolecules comprising a lipid core of triglycerides and cholesteryl esters that is surrounded by a hydrophilic phospholipid monolayer. The latter is decorated with apolipoproteins such as ApoB and ApoE that stabilize the complex and provide a functional identity (6). ApoE is synthesized primarily in hepatocytes and several non-hepatic tissues, including the brain, artery walls, spleen, kidney, muscle, and adipose tissue, but most LP subclasses in the plasma associate with hepatocyte-derived ApoE (6–8). ApoE regulates the clearance of cholesterol-rich LPs from circulatory systems via its binding to receptors on the surface of hepatocytes, including heparan sulfate proteoglycan (HSPG) and low-density lipoprotein receptor (LDLR) (6). It was reported that liver-generated ApoE is superior to ApoE from other tissues in the clearance of LP remnants (9). Abnormal function of ApoE was found in patients with type III hyperlipoproteinemia, which is a disorder characterized by high blood levels of triglycerides and cholesterol (10, 11). Moreover, a recent study reported that liver-generated ApoE affects integrity of the brain (12). At least 18 diseases, including Alzheimer’s and cardiovascular diseases are strongly associated with *APOE* genotypes (13). Moreover, *APOE* genotypes appear to correlate with the outcome of some viral diseases such as coronavirus disease of 2019 (COVID-19) (14–17).

Notably, hepatic ApoE is an essential integral component of HCV and hepatitis B virus (HBV) particles and has been suggested to be a promising target for the development of effective HCV vaccines (18, 19). In addition, recent findings indicate that plasma ApoE and other apolipoproteins form a protein coat around secreted extracellular vesicles (EVs) and affect EV signaling function (20–22).

Despite a long history of intensive research, the trafficking, egress, and transmission route of hepatic ApoE-LPs are poorly understood. ApoE is a 299 amino acids (aa) long protein that contains an 18 aa N-terminal signal peptide targeting the protein co-translationally into the ER lumen (23, 24). ER-luminal ApoE is transported to the Golgi where it is modified by O-linked sialylation (25, 26) and associates with nascent LPs containing ApoB100 and triglycerides. Thereafter, ApoE-ApoB100-containing LPs are further lipidated giving rise to mature LPs that have lower buoyant density (6, 25–27). To be secreted, mature LPs must be transported from the *trans*-Golgi network (TGN) to the plasma membrane (PM), but this process is poorly understood. By using an *in vitro* assay, Hossain and colleagues reported a novel transport vesicle delivering VLDL to the PM of rat hepatocytes, but the identity of this vesicle class is unknown (28). At least in macrophages, secretion of ApoE follows the microtubule network along a protein kinase A and calcium-dependent pathway (29). In addition, in a pigment cell type ApoE associates with intraluminal vesicles (ILVs) within endosomes and is released with these vesicles in the form of exosomes (30). The observation that inhibition of ApoE sorting to endosomes retains ApoE at the Golgi compartment argues for Golgi–endosome transport of ApoE (30). The endosomal compartment is also required for the export of HCV particles that are thought to follow a noncanonical secretory route (31). Since HCV particles associate intracellularly with hepatic ApoE (17, 32) hepatocyte-derived ApoE-LPs might also be released via an endosomal egress pathway. Consistently, HCV hijacks the endosomal pathway for the transmission of viral RNA genomes via endosome-derived CD63-positive extracellular vesicles (EVs) (33–36).

To study the egress pathway of ApoE-LPs, we established a fully functional fluorescently tagged ApoE and show that ApoE-LPs enrich in CD63-positive endosomes of hepatocytes. Intracellular ApoE-LPs and CD63-positive EV precursors associate with each other and are co-secreted for cell-to-cell transmission. Expanding these observations to HCV, we report that the viral replicase factor nonstructural protein 5A (NS5A) associates with ApoE. This interaction is required for the release of ApoE-associated CD63-positive EVs containing viral RNA and being taken up by non-infected bystander cells. Thus, endosomal release of ApoE-LPs appears to be a physiological pathway that is exploited by HCV.

## Results

### Establishment of fully functional fluorescently tagged ApoE

Live-cell imaging of ApoE requires a suitable fluorescently tagged protein that retains full functionality. GFP was previously selected for ApoE labeling, but ApoE-GFP fusion proteins are prone to undesired cleavage and lack full functionality (37). To overcome this limitation, we probed alternative fluorescent protein (FP) tags that were fused to the C-terminus of ApoE. As target cells, we employed hepatocyte (Huh7)-derived cells (cell line Huh7-Lunet/CD81H), because they are well suitable for various imaging approaches (32) and highly permissive to HCV (38). To avoid excessive overexpression, endogenous ApoE amount was reduced to undetectable level by stable knockdown, prior to lentiviral transduction of the cells with constructs encoding various ApoE fusion proteins (FPs) (32). Western blot analysis revealed that in the case of all ApoE-redFPs, in addition to the full-length proteins (∼58-kDa), truncated proteins (∼46 kDa) were detected (Fig S1A). This truncation might be due to hydrolysis of the N-acylimine group of the DsRed-like chromophores in these FPs, especially under the acidic conditions in late endosomes where ApoE is expected to reside. Therefore, we tagged ApoE with mTurquoise2 (mT2) and eYFP. Consistent with our assumption, ApoE^mT2^ and ApoE^eYFP^ were not fragmented (Fig S1A, upper right). Because mT2 is a rapidly-maturing cyan monomer with very low acid sensitivity (pKa = 3.1) (39), we selected this tagged ApoE for functional validation.

ApoE^mT2^ was efficiently secreted into the cell culture supernatant (Fig 1A). Moreover, the association of secreted LPs with ApoE^mT2^ was well comparable to the one with wildtype (wt) ApoE as determined by separation of LPs using sucrose density gradient centrifugation (peak density of ApoE^mT2^ and ApoE^wt^ = 1.05 vs. 1.04 g/ml, respectively) (Fig 1B). Moreover, in addition to a weak and diffuse ER-like pattern, ApoE^mT2^ formed strong and dotted puncta characteristic for LPs and colocalized with ApoB, a well-established LP marker (Fig 1C). Of note, ApoE puncta detected by immunofluorescence in fixed cells were much dimmer than those containing mT2, thus increasing sensitivity of our analyses, especially in live-cell imaging (Fig 1C). We further investigated ApoE^mT2^ subcellular distribution in nonhepatic cells having undetectable levels of ApoE such as HEK293T and Hela cells (30, 40). Upon ectopic expression of ApoE^mT2^, we observed a dot-like pattern in both cell lines, which was well comparable to the one detected in Huh7-Lunet cells (Fig S1B).

**Fig 1.**
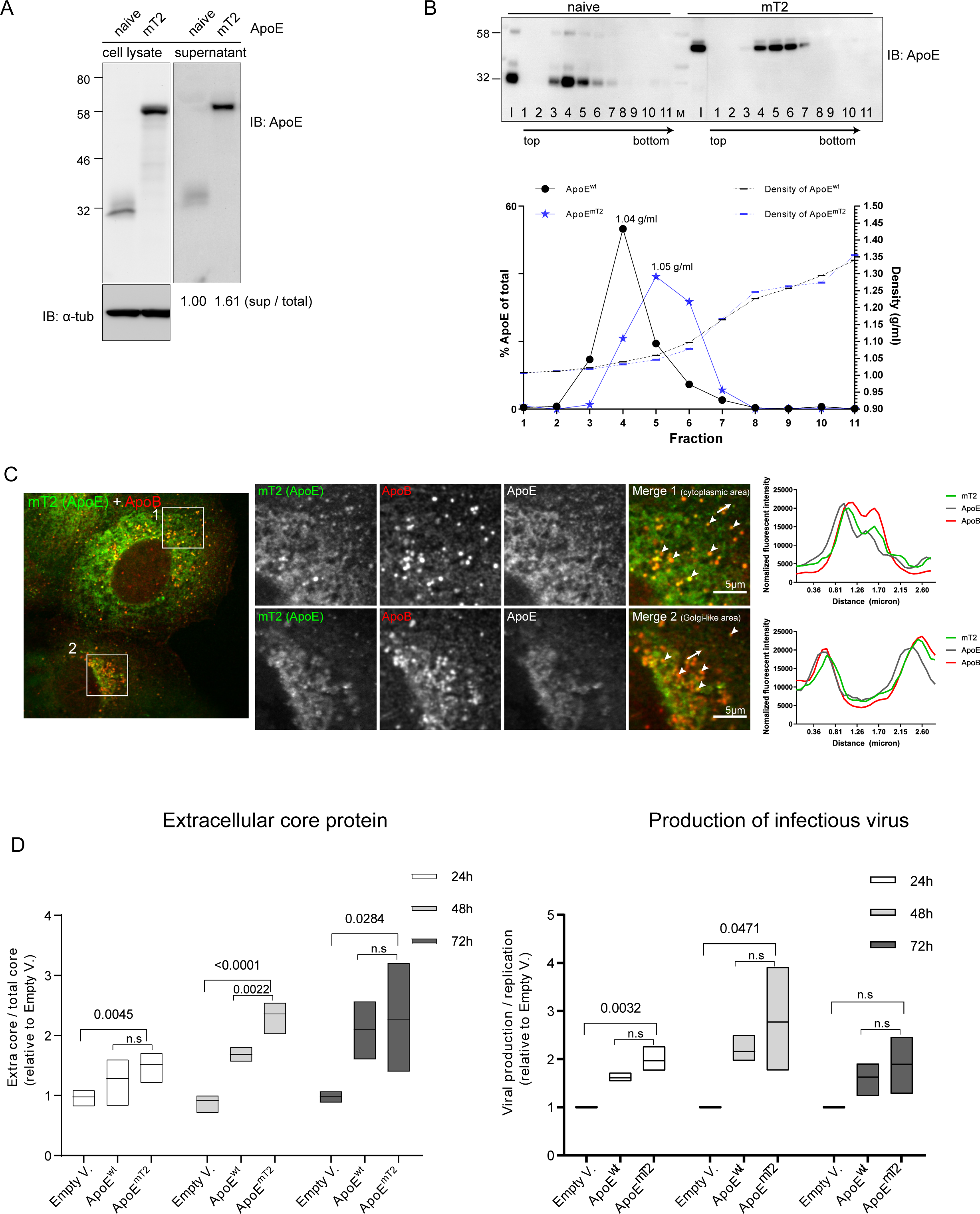
Establishment and characterization of fully functional fluorescently tagged ApoE. (A) Secretion of ApoE^mT2^. Lysates and supernatants of Huh7-Lunet/ApoE-KD cells stably expressing or not mTurquoise2-tagged ApoE were harvested one day after seeding and samples were analyzed by Western blot using ApoE-specific antibody; α-tubulin served as a loading control for cell lysates. The ratios of secreted to total ApoE are given below the lanes. The value of ApoE^wt^ was set to 1. (B) Density of secreted ApoE^mT2^. Upper panel: conditioned media of naïve ApoE and ApoE^mT2^-reconstituted cells from (A) were subjected to 10-50% iodixanol isopycnic centrifugation and fractions were analyzed by Western blot using ApoE-specific antibody. I: input, M: protein marker lane. Lower panel: signal intensities of the Western blot image were quantified and values were normalized to total ApoE amount in all fractions. Densities of fractions are specified on the right Y-axis (g/ml). Densities of peak fractions are given. (C) Normal lipid-binding property of ApoE^mT2^. Immunofluorescent staining of ApoE^mT2^ in ApoE^mT2^ reconstituted Huh7-Lunet/ApoE-KD cells using ApoE- and ApoB-specific antibodies. Two-row images on the right show magnified views of boxed areas in the left overview image. Arrowheads in cropped images point to signal overlaps of ApoE^mT2^ and ApoB; plot profiles in the right panels are along the lines indicated with white arrows in the merge images. (D) Functionality of ApoE^mT2^ as determined by the rescue of infectious HCV particle production. Left panel: Huh7-Lunet/ApoE-KD cells were transduced with either an empty vector (Empty V.), or ApoE^wt^, or ApoE^mT2^ and stably selected. Cells were then electroporated with *in vitro* transcripts of the Renilla luciferase (RLU) HCV reporter genome JcR2a. At 24, 48 and 72 h post-electroporation, amounts of extracellular core protein present in culture supernatants were determined by chemiluminescence assay. Right panel: amount of infectious HCV particles released into the culture supernatant of electroporated cells. At the indicated time points supernatants were harvested, naïve Huh7.5 cells were inoculated and 72 h later, luciferase activity was determined. Values were normalized to HCV RNA replication in each cell line to exclude replication effects. Data are medians (range) from three independent experiments. P-value was determined using unpaired Student’s *t*-test. N.s: not statistically significant (P>0.05).

Next, we validated the functionality of ApoE^mT2^ by probing its capacity to rescue the production of infectious HCV, which was used as readout because this virus incorporates ApoE into virions intracellularly to increase viral infectivity (32, 41).To facilitate the analysis, we employed the HCV reporter virus JcR2a encoding Renilla luciferase (42). JcR2a *in vitro* transcripts were transfected into ApoE knock-down Huh7-Lunet cells expressing ApoE^mT2^ or ApoE^wt^ or containing the empty expression vector. RNA replication, determined by luciferase assay and intracellular accumulation of core protein, was comparable among all 3 cell pools (Figs S1C and S1D). As expected, ApoE^wt^ and ApoE^mT2^ expression significantly alleviated the secretion of HCV virions as determined by quantifying extracellular HCV core protein and infectivity assay (Fig 1D). Baseline production of HCV in empty vector-transduced Huh7-Lunet cells was further reduced when we used the nonhepatic cell line HEK293T-miR122, which does not express endogenous apolipoproteins but supports HCV RNA replication (40), arguing that the expression of non-ApoE LPs in Huh7-derived cells compensates, at least in part, for ApoE deficiency (43, 44) (Fig S2). Also in these cells, ApoE^wt^ and ApoE^mT2^ rescued infectious HCV particle production (Fig S2). Taken together, our data show that mT2 is a novel and well-applicable tag for labeling and functional analyses of ApoE.

### Endosomal trafficking and egress of ApoE in hepatocytes

Having established a suitable FP-tagged ApoE, we employed light and electron microscopy methods to study the trafficking and egress route of ApoE in hepatocytes. First, we confirmed the conventional trafficking route of ApoE, which starts at the ER where it is co-translationally delivered into the lumen to enter the secretory pathway (23, 24). Consistently, in Huh7-Lunet/ApoE^mT2^ cells we detected reticular ApoE^mT2^ signals overlapping with the ER marker PDI (Fig 2A, top row). In addition, we observed condensed ApoE^mT2^ puncta in the Golgi area containing GM130, a marker of the Golgi apparatus, consistent with the assembly of ApoE-LPs at this site (Fig 2A, middle row).

**Fig 2.**
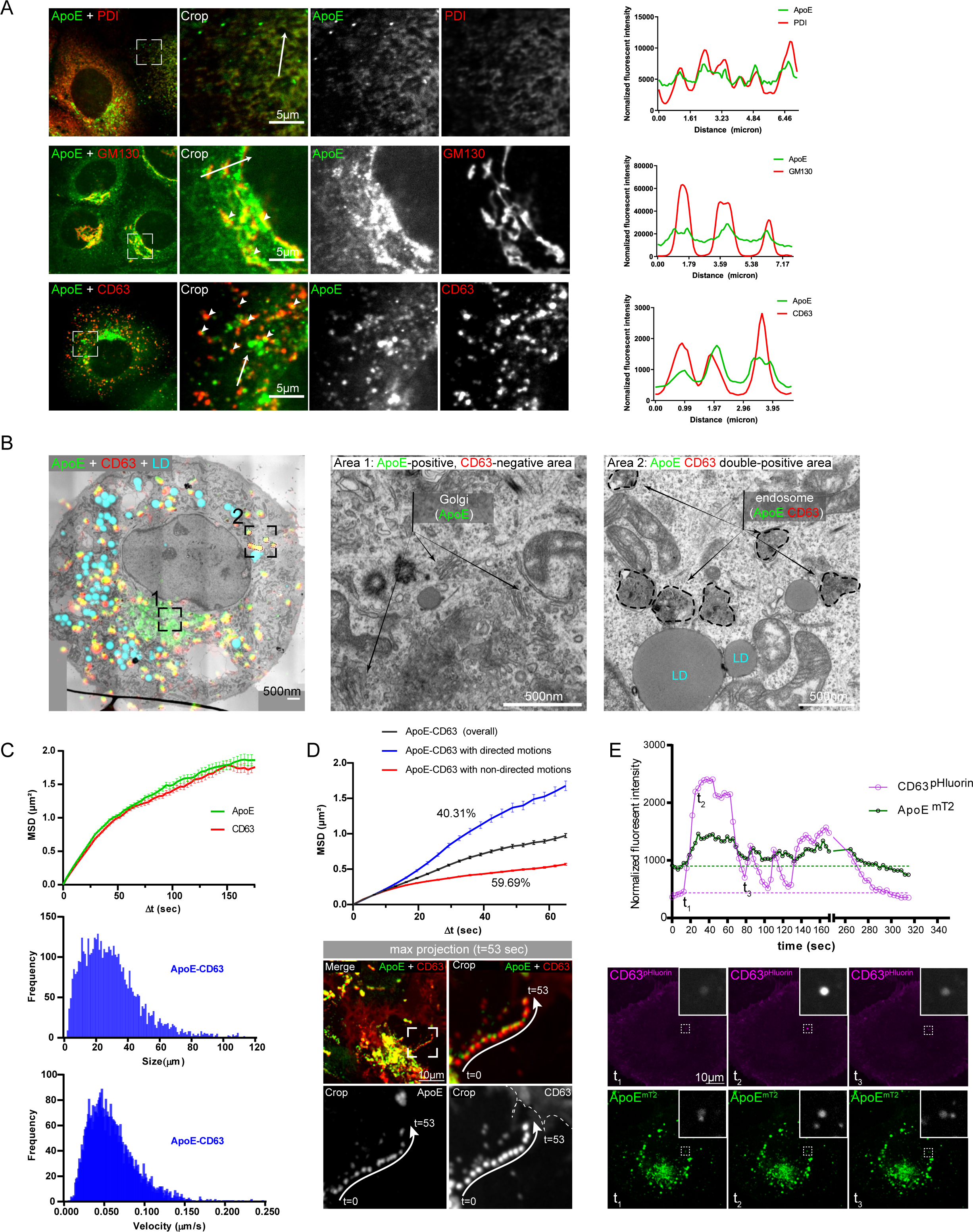
Detection of ApoE in CD63-positive late endosomes, intracellular endosomal trafficking of ApoE and egress from hepatocytes. (A) Colocalization of ApoE^mT2^ with markers of the ER (PDI), Golgi (GM130), and intraluminal vesicles/endosomes (CD63). Proteins specified on the top of each panel were detected in Huh7-Lunet/ApoE^mT2^ cells by immunostaining and cells were analyzed by confocal microscopy. Profiles on the right of each panel were taken along the lines indicated with white arrows in cropped images. (B) Endosomal localization of ApoE-CD63 double-positive structures. Huh7-Lunet/ApoE^mT2^ cells expressing CD63^mCherry^ were analyzed by CLEM using lipid droplets (LDs) stained with lipidTox as fiducial markers. The overlay image is shown on the left. Middle and right panels: magnified EM micrographs from an area with ApoE-positive, CD63-negative signals showing Golgi stacks and vesicles (crop 1) and from an area with ApoE-CD63 double-positive endosomes (crop 2), respectively. (C-E) Secretion of ApoE-associated CD63-positive EVs. (C) Motility of intracellular ApoE-CD63 double-positive structures. [Top] Mean squared displacement (MSD) of general ApoE and CD63 trafficking. [Middle] Sizes of ApoE-CD63 double-positive structures. [Bottom] Trafficking velocities of ApoE-CD63 double-positive structures. (D) [Top] Mean squared displacement (MSD) of general ApoE and CD63 trafficking and those with directed and non-directed motions. [Bottom] Example of ApoE-CD63 co-trafficking by a directed motion. Huh7-Lunet/ApoE^mT2^ cells expressing CD63^mcherry^ were analyzed by live-cell confocal microscopy. A maximum projection image showing co-trafficking of an ApoE-CD63 complex with a directed motion to the cell periphery is shown. Frame interval = 2.65 sec; whole duration = 53 sec. (E) Secretion of ApoE-positive ILVs visualized by pHluorin-tagged CD63. Huh7-Lunet cells expressing ApoE^mT2^ and CD63^pHluorin^ were cultured in imaging medium (pH 7.4) and analyzed by time-lapse confocal microscopy with a focus on plasma membrane resident CD63-fluorescent signals. [Top] Maximum fluorescence intensity of CD63^pHluorin^ and associated ApoE in the selected dashed area indicated in supplementary movie 2. Images taken at indicated time points are displayed on the bottom and they correspond to initiation (t1), peak (t2), and late-secretion (t3), respectively.

To determine whether hepatocyte-derived ApoE-LPs are released via an endosomal egress pathway, we initially determined its colocalization with CD63, the commonly used marker of ILVs that are sorted into late endosomes (45, 46). We detected numerous ApoE^mT2^-containing structures in Golgi-devoid areas and these signals predominantly overlapped with CD63, indicating accumulation of ApoE in late endosomes (Fig 2A, bottom row). Consistently, a fraction of ApoE signals overlapped strongly with Rab7 (a marker of late endosomes), but rarely with ADRP (a marker of lipid droplets) (Fig S3). We further identified the ultrastructure of ApoE-CD63 positive signals by correlative light and electron microscopy (CLEM) using lipid droplets as fiducial markers, because they are easy to detect in both light and electron microcopy and have a unique distribution and size in each Huh7 cell (Fig 2B). We found that ApoE-CD63 double-positive signals predominantly corresponded to regions containing electron-dense vesicles of ∼500 nm in diameter, which is a typical feature of endosomal compartments (47) (Fig 2B, right panel).

With the aim to track and record the dynamics of ApoE association with CD63, we took advantage of ApoE^mT2^ and conducted time-lapse confocal microscopy (Movie S1). We observed co-trafficking of ApoE-CD63 double-positive puncta as indicated by their similar mean squared displacement values (Fig 2C, top). Particle size and velocity of ApoE-CD63 double-positive signals were also computed, revealing substantial heterogeneity of particle motions (Fig 2C, middle and bottom). Importantly, a subpopulation of these vesicles displayed directed motions (Fig 2D, upper panel), suggesting microtubule-dependent trafficking of late endosomes containing ApoE and CD63 (48, 49). An example of ApoE-CD63 co-trafficking dynamics showing a directed motion towards the cell periphery is shown in Fig 2D, lower panel.

To visualize the intracellular trafficking of ApoE-associated CD63-positive ILVs, we took advantage of the acidic pH in endosomes that gets neutral as endosomes fuse with the PM to release ILVs contained therein. As an endosome-PM fusion sensor, we employed an improved version of pHluorin (50) that was inserted into the first extracellular loop of CD63, thus exposing pHluorin to the acidic environment of the endosomes. The signal of pHluorin-tagged CD63 is quenched in the endosomes and is exclusively excited upon exposure of the endosomes’ interior to the neutral pH of the extracellular environment (51). To capture ApoE-CD63 co-secretion, we used time-lapse live-cell confocal microscopy by setting the focal plane to the PM as determined by the basal fluorescence of the CD63^pHluorin^ signal (Fig 2E and Movie S2). As expected, CD63^pHluorin^ expressed in Huh7-Lunet cells showed a predominant fluorescent signal in the PM. Of note, we observed occasional steep and rapid increases of the vesicular ApoE-associated pHluorin signal (Fig 2E, time point t_2_) corresponding most likely to the fusion of ApoE-CD63 containing endosomes with the PM and thus, the release of ApoE-associated CD63-positive ILVs.

### Co-secretion and cell-to-cell co-transmission of ApoE and endosome-derived extracellular vesicles

A recent study by Busatto and colleagues demonstrated that EVs in crude plasma frequently bind to and fuse with LPs arguing for a physiological interaction between these two nano-particle species (20). Given the cotrafficking of intracellular hepatic ApoE with CD63 and the secretion of ApoE-associated CD63 (Fig 2), we speculated that extracellular hepatic ApoE might associate with CD63-positive EVs via LPs. Given the difficulties to separate EVs from LPs (21, 52–54), we employed ApoE-specific pull-down to isolate ApoE from the supernatant of Huh7-Lunet cells that had been cultured in EV-depleted medium. Captured complexes were eluted under native conditions and analyzed by EM revealing predominantly small vesicles, which had the size of regular LDL or large HDL particles (mean diameter ∼25 nm) (Fig 3A). Of note, we detected in much lower quantity co-captured bigger vesicles (mean diameter ≥50 nm) (Fig 3A, labeled with stars), a fraction of them staining positive for CD63 and being associated with the smaller ApoE-positive particles (Fig 3B). This result argued for the stable interaction between secreted ApoE-LPs and CD63-positive EVs, consistent with a previous study (20).

**Fig 3.**
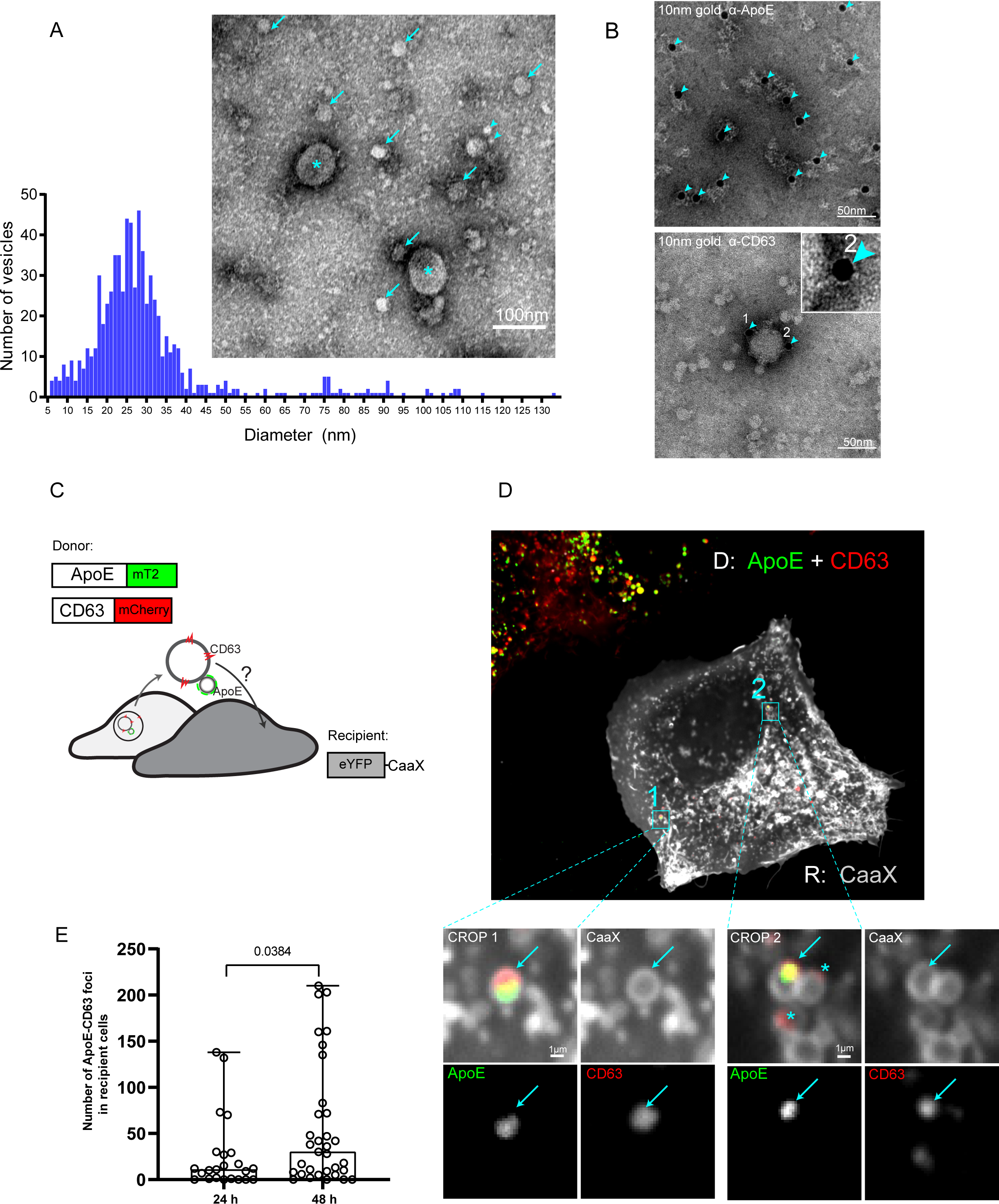
Co-secretion and cell-to-cell co-transmission of ApoE with endosome-derived extracellular vesicles. (A) Visualization of ApoE-containing EVs. Huh7-Lunet cells were cultured in EV-depleted medium and ApoE-associated vesicles released into the culture medium were captured using ApoE-specific antibody. Immunocomplexes were analyzed by TEM after negative staining. Arrowheads: ∼5-10 nm vesicles; arrows: ∼20-30 nm vesicles; stars: ∼50-60 nm vesicles. Vesicles in the electron micrographs were segmented by using Ilastik to allow quantification of vesicle diameters shown in the histogram below the micrograph. (B) Association of secreted lipoproteins with EVs. Purified ApoE-associated vesicles from (A) were immunogold-labeled with ApoE-(upper) and CD63-specific antibodies (lower). Arrowheads point towards gold particles. A zoom image of a CD63-positive gold particle is shown on the top. Note that immuno-gold labeling of ApoE alters the vesicular shape of LPs, most likely because of distortion of ApoE during the labeling procedure thereby destabilizing the LP structure. (C-E) Visualization of the co-uptake of LP-EV complexes by recipient cells. (C) Schematic representation of used approach. Huh7-Lunet/ApoE^mT2^/CD63^mCherry^ served as donor cells; Huh7-Lunet cells expressing eYFP-tagged CaaX (the farnesylation signal from human HRAS) as recipients. (D) Donor and recipient cells from (C) were co-cultured for 16 h and analyzed by live-cell confocal imaging (refers to supplementary movie 3). D: donor; R: recipient. Arrows in cropped sections on the bottom indicate transferred ApoE-CD63 signals; stars: transferred CD63-only signals. (E) Donor and recipient cells from (C) were co-cultured and fixed at 24 h and 48 h post-seeding. The numbers of ApoE-CD63 double-positive signals in single recipient cells were quantified. Each dot represents a single cell. P-value was determined using unpaired Student’s *t*-test.

Next, we examined the possible cell-to-cell co-transfer of ApoE-LPs associated with CD63-positive EVs. To this end, we used hepatic donor cells expressing fluorescently labeled ApoE^mT2^ and CD63^mCherry^, and recipient cells expressing the human HRAS-derived CaaX peptide that was fused to eYFP (Fig 3C, D, gray cells). In this fusion protein the CaaX motif, which is a farnesylation signal, targets the protein to cellular membranes making them easily trackable via eYFP and allowing the faithful discrimination of recipient and donor cells. Cells were seeded into imaging dishes and 16 h later, examined by live-cell confocal microscopy. We observed donor-derived ApoE^mT2^-CD63^mCherry^ double-positive structures in recipient cells, indicating transfer and uptake of these structures (Fig 3D and Movie S3). Quantitative image analysis revealed a time-dependent increase in the number of ApoE^mT2^-CD63^mCherry^ double-positive structures in single recipient cells, especially at 48 h post-seeding (Fig 3E). Taken together, this result indicates intercellular transmission of endosome-derived EVs bound to hepatic ApoE-LPs.

### Intracellular enrichment of HCV NS5A in ApoE-positive structures independent from virion assembly

As alluded to in the introduction, ApoE associates with HCV particles, most likely via the viral envelope glycoprotein complex E1/E2 (32, 55) and with the viral replicase factor NS5A (56–58). While the ApoE-E1/E2 interaction appears to be critical for HCV particle production, NS5A has been detected in purified EV preparations (59, 60), raising the question of whether NS5A follows the ApoE endosomal egress pathway. To address this question, we monitored ApoE, NS5A and E2 trafficking in HCV-replicating Huh7-Lunet cells stably expressing ApoE^mT2^. FPs for NS5A and E2 were chosen to allow clear spectral separation from each other and from ApoE^mT2^. In each case, fusion with the FP did not affect the functionality of the protein as shown here for ApoE^mT2^, and earlier for tagged NS5A and E2 (61, 62). To allow live-cell imaging under low biosafety conditions, we took advantage of the HCV trans-complementation system (63) in which the HCV genome is genetically split into a stably expressed unit encoding the viral assembly factors (core-E1-E2^eYFP^-p7-NS2) and a self-replicating subgenomic replicon encoding the viral replicase proteins (NS3-4A-4B-5A^mCherry^-5B) (Fig 4A). To determine the overall subcellular distribution of FP-tagged ApoE^mT2^, NS5A^mCherry^, and E2^eYFP^ during the course of HCV infection, we acquired time-lapse images by confocal spinning disc microscopy in 30 min intervals between 5 and 54 h post-electroporation using minimum laser exposure to avoid phototoxicity. Prior to electroporation of the subgenomic replicon, E2^eYFP^ showed a reticular ER-like pattern consistent with its ER retention (64). Around 25 h post-electroporation, E2^eYFP^ subcellular distribution began to change and NS5A^mCherry^-E2^eYFP^ double-positive foci became visible (Fig 4B, arrowheads; Movie S4) (62). In addition, ApoE^mT2^-

**Fig 4.**
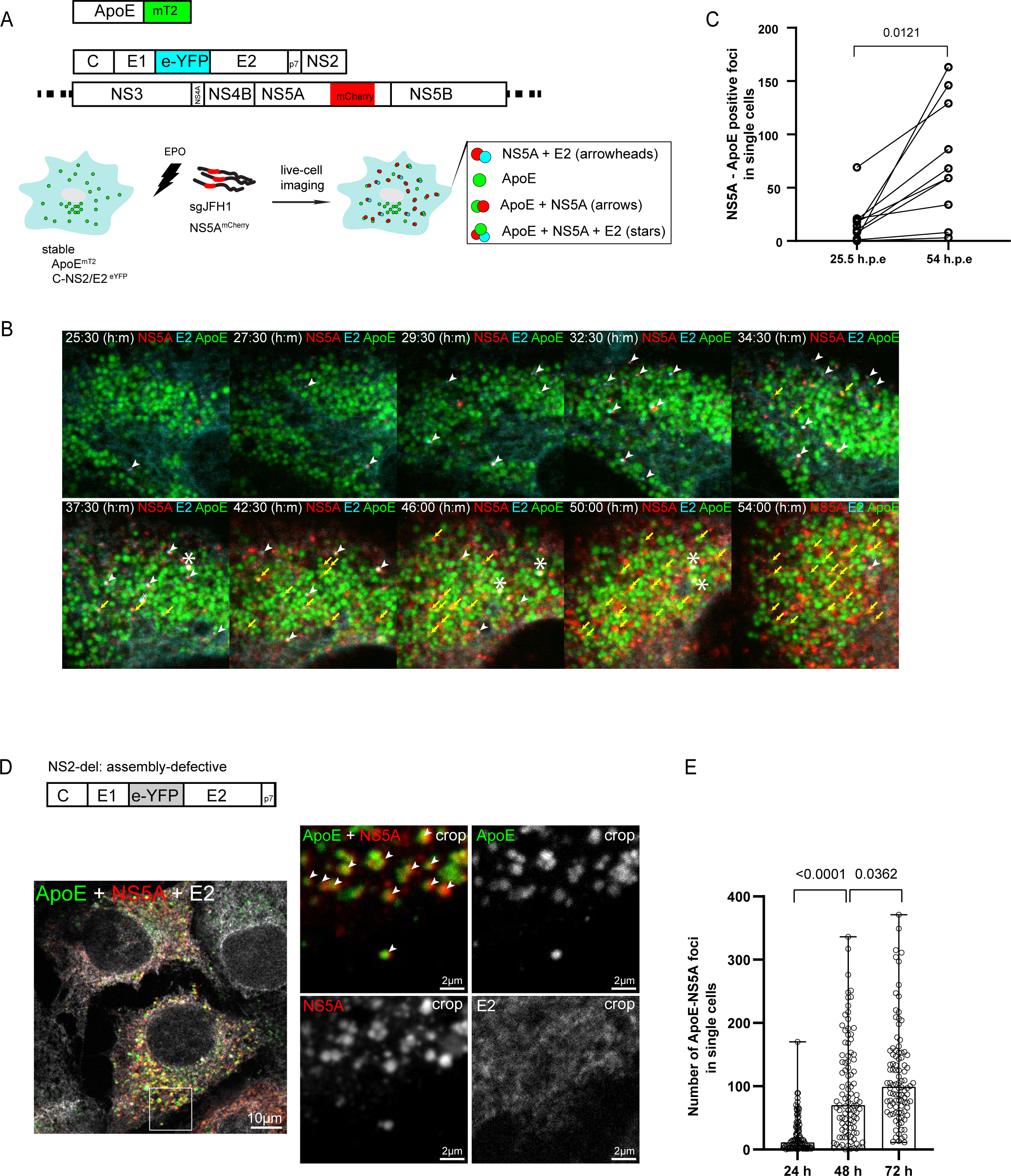
Enrichment of NS5A in ApoE-positive structures and co-trafficking of ApoE^mT2^ with NS5A and E2 independent of HCV assembly. (A) Experimental approach. Fluorescently tagged ApoE^mT2^, HCV proteins supporting assembly (C to NS2 with eYFP-tagged E2), and a subgenomic replicon (dotted lines indicate 5’ and 3’ NTRs) are shown from top to bottom; the experimental approach is depicted below. Cells stably expressing ApoE^mT2^ and C-NS2/E2^eYFP^ were electroporated with the replicon RNA encoding mCherry-tagged NS5A. Cells were subjected to confocal time-lapse live-cell imaging to monitor signal overlaps of the various fluorescent proteins: NS5A + E2; ApoE only; ApoE + NS5A; ApoE + NS5A + E2. (B) Time-dependent enrichment of NS5A-ApoE double-positive structures in HCV-replicating cells. Huh7-Lunet/ApoE^mT2^ cells expressing HCV Core-NS2/E2^eYFP^ and containing the subgenomic replicon were subjected to live-cell confocal imaging from 5 to 54 h p.e (30 min/frame) to observe ApoE, NS5A, and E2 signals. A series of still images taken at time points after electroporation specified on the top are shown. White arrowheads: NS5A-E2 foci; yellow arrows: ApoE-NS5A foci; stars: ApoE-NS5A-E2 triple-positive foci. (C) Quantification of NS5A-ApoE double-positive foci detected in single cells in (B). Ten single cells were analyzed. P-value was determined using Mann-Whitney test. (D) Assembly-independent enrichment of NS5A in ApoE-positive foci. Huh7-Lunet/ApoE^mT2^ cells expressing the C-p7 construct (NS2-deletion; upper panel) were electroporated with *in vitro*-transcripts of the subgenomic replicon sgJFH1/NS5A^mCherry^ and analyzed by confocal microscopy to observe ApoE, NS5A, and E2 signals. A representative image showing ApoE-NS5A double-positive foci (arrowheads) and diffuse E2 signal at 72 h p.e is shown. Images on the right show magnified views of the boxed area in the left overview image. (E) Quantification of NS5A-ApoE double-positive foci detected in 100 single cells in (D) at 24, 48, and 72 h p.e. Data are medians (range) of the number of detected foci. P-value was determined using unpaired Student’s *t*-test.

NS5A^mCherry^-E2^eYFP^ triple-positive foci, putative sites of HCV assembly, were observed, but their abundance was very low (Fig 4B, stars). Consistent with ongoing HCV replication, NS5A^mCherry^ signal intensity increased steadily and NS5A^mCherry^-ApoE^mT2^ double-positive foci formed. Their abundance increased significantly over time (Fig 4C), much higher as compared to NS5A^mCherry^-E2^eYFP^ positive foci. We confirmed the high number of NS5A^mCherry^-ApoE^mT2^ double-positive foci at a late stage of infection by live-cell imaging using a shorter time interval (10 sec/frame). Under this imaging condition, NS5A^mCherry^-ApoE^mT2^ foci were readily detectable (Movie S5).

To confirm the formation of NS5A-ApoE double-positive structures in the context of a full-length HCV genome, we transfected Huh7-Lunet/ApoE^mT2^ cells with *in vitro* transcripts of a cloned HCV genome and determined NS5A and ApoE subcellular distribution in relation to the ER marker PDI by immunofluorescence. Also under these conditions, ApoE signals significantly overlapped with NS5A, confirming that the trans-complementation system faithfully recapitulates events occurring in natural infection (Figs S4A and S4B).

Next we determined whether formation of NS5A-ApoE positive structures depends on viral assembly or is linked to some other events such as the formation of EVs. To this end, we used the same experimental approach as shown in Fig 4A, but employed a construct lacking the viral assembly factor NS2 (core-E1-E2^eYFP^-p7) (Fig 4D, upper panel) (65). While under these conditions NS5A^mCherry^-E2^eYFP^ double-positive structures were no longer detected, NS5A^mCherry^-ApoE^mT2^ double-positive dots still formed (Figs 4D and 4E). These results suggested that enrichment of NS5A in ApoE-positive puncta does not depend on HCV assembly.

### Formation of NS5A- and ApoE-containing intraluminal vesicles in CD63-positive endosomes

Since HCV has been reported to transmit its RNA via a noncanonical pathway comprising endosome-derived CD63-positive EVs (34–36, 66) and because ApoE-LPs also egress along the CD63-positive late endosomal pathway (Figs 2 and 3), we characterized the association of ApoE-NS5A double-positive structures with CD63 in greater detail by using super-resolution microscopy. To make this possible, we exchanged the FPs of ApoE^mT2^ and NS5A^mCherry^ for SNAPf and CLIPf, respectively (Fig 5A, upper). Both fusion proteins were fully functional as revealed by the secretion of ApoE^SNAPf^ and the replication competence of NS5A^CLIPf^ (Figs S4C and S4D, respectively). In the first set of experiments, Huh7-Lunet/ApoE^SNAPf^ cells were transfected with subgenomic replicon RNA and 48 h later, cells were incubated with medium containing the dyes SNAP-SIR647 and CLIP-ATTO590, respectively for 1 h. Thereafter, cells were washed to remove unbound dyes and subjected to live-cell imaging or fixed-cell microscopy (Fig 5A, lower). Confocal imaging of the cells revealed specific labeling of ApoE^SNAPf^ and NS5A^CLIPf^ and strong colocalization of both proteins (Fig 5B), consistent with our previous results with FP-tagged ApoE and NS5A (Fig 4). Importantly, we found that about half of ApoE - NS5A double-positive foci also contained CD63 (Fig 5C, Fig S4E). When we visualized NS5A and ApoE by super-resolution STED microscopy, in addition to the reticular ER and the ring-like lipid droplet staining patterns of NS5A^CLIPf^, we detected ∼100-200 nm diameter NS5A^CLIPf^-positive structures that were decorated with ApoE^SNAPf^ at CD63-positive sites (Fig 5D, arrows).

**Fig 5.**
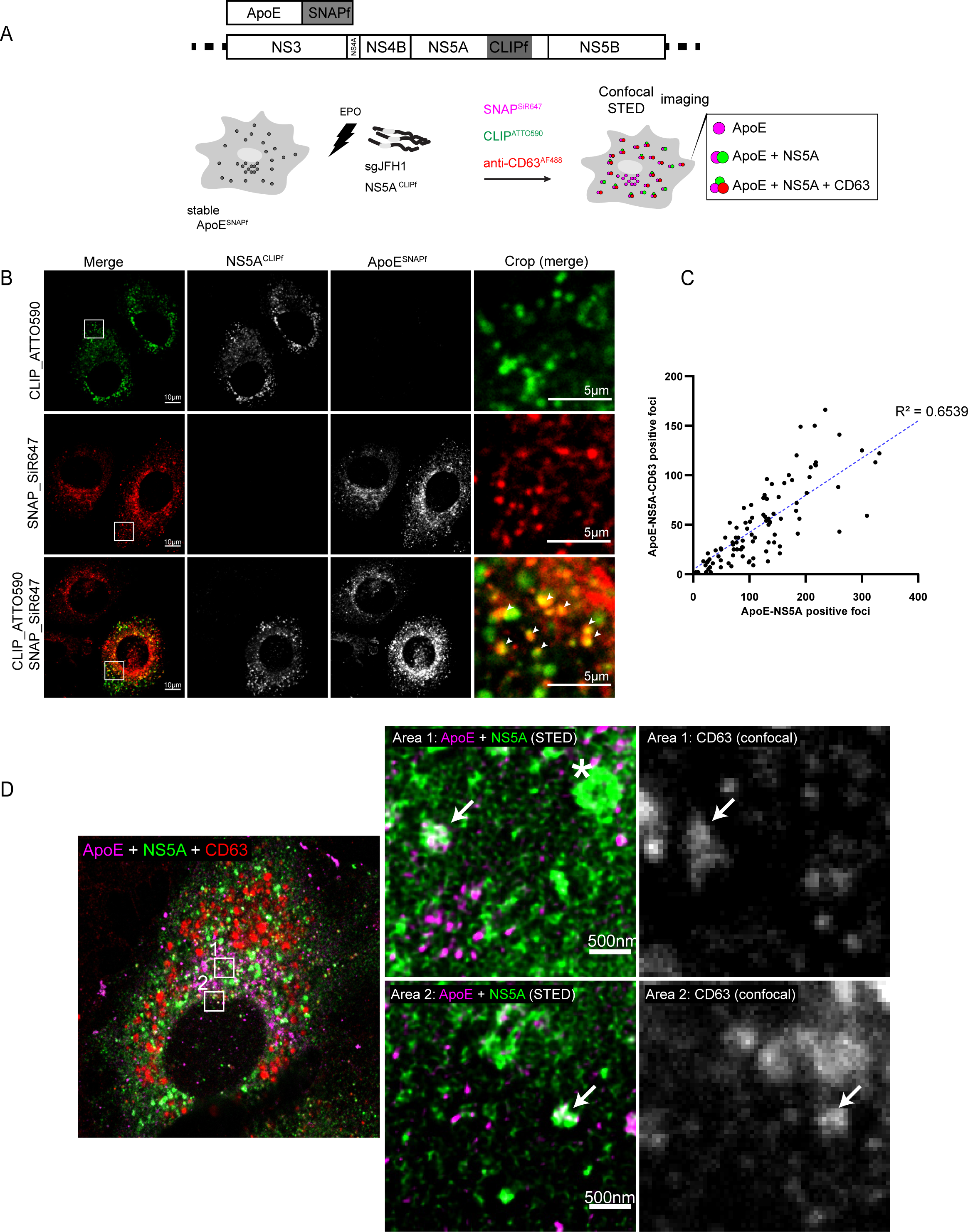
Colocalization of NS5A and ApoE with the intraluminal vesicle marker CD63 as revealed by super resolution microscopy. (A) Experimental approach. Schematic representations of SNAPf-tagged ApoE and the subgenomic replicon encoding CLIPf-tagged NS5A are shown on the top. Huh7-Lunet cells were lentivirally transduced with the ApoE expression vector and transfected with the subgenomic replicon RNA. ApoE and NS5A were detected by STED microscopy and CD63 by immunofluorescence confocal microscopy. (B) Colocalization of ApoE^SNAPf^ and NS5A^CLIPf^. Huh7-Lunet/ApoE^SNAPf^ cells were electroporated with subgenomic replicon RNA encoding NS5A^CLIPf^ and after 48 h, cells were labeled with SNAP^SiR647^ and CLIP^ATTO590^ for 1 h, fixed, and subjected to confocal microscopy. Arrowheads: colocalized ApoE-NS5A signals. (C) Quantification of CD63-positive ApoE-NS5A double-positive foci. Cells from (B) harvested 72 h p.e were fixed, permeabilized, and incubated with anti-CD63^AF488^ antibody. To determine the correlation between ApoE-NS5A double-positive foci and how many of them colocalized with CD63, 100 cells were analyzed. Each dot represents one cell and displays the number of ApoE-NS5A double-positive foci (x-axis) and the number of CD63-ApoE-NS5A triple-positive foci (y-axis). The R-squared value is given on the plot. (D) STED-resolved ApoE-NS5A double-positive structures colocalizing with the intraluminal vesicle marker CD63. Huh7-Lunet/ApoE^SNAPf^ cells were electroporated with the subgenomic replicon RNA encoding NS5A^CLIPf^. After 48 h, cells were labeled with SNAP^SiR647^ and CLIP^ATTO590^ for 1 h, fixed, and incubated with anti-CD63^AF488^ antibody. ApoE, NS5A, and CD63 fluorescent signals were sequentially imaged using confocal and STED microscopy, the latter to achieve a higher resolution of ApoE and NS5A signals that were deconvoluted using Huygens. Arrows: ∼100-200 nm-sized ApoE-NS5A-CD63 positive structures; star: ∼500 nm-sized ring-like NS5A positive structure.

To determine the ultrastructure of ApoE-NS5A double-positive sites, we employed CLEM using Huh7-Lunet/ApoE^mT2^ cells expressing the HCV assembly factors and containing a subgenomic replicon (refer to Fig 4A). We observed an overlap of NS5A^mCherry^-ApoE^mT2^ double-positive signals with endosomes (Figs 6A and 6B). Strikingly, inside these endosomes, we detected numerous ILVs with double or multi-membrane bilayers (Fig 6B, crop 1, 2, and 3, arrowheads), which were only rarely detected in NS5A^mCherry^-negative, ApoE^mT2^-positive endosomes (crop 4). Sites of NS5A^mCherry^-E2^eYFP^ double-positive structures, putative HCV assembly sites, overlapped with HCV-induced accumulations of double-membrane vesicles (DMVs), the presumed sites of viral RNA replication that were often found in close proximity to lipid droplets (Fig 6B, crop 5 and 6) as reported earlier (62). Taken together, these results argued for the accumulation of NS5A- and ApoE-containing ILVs at sites of CD63-positive endosomes.

**Fig 6.**
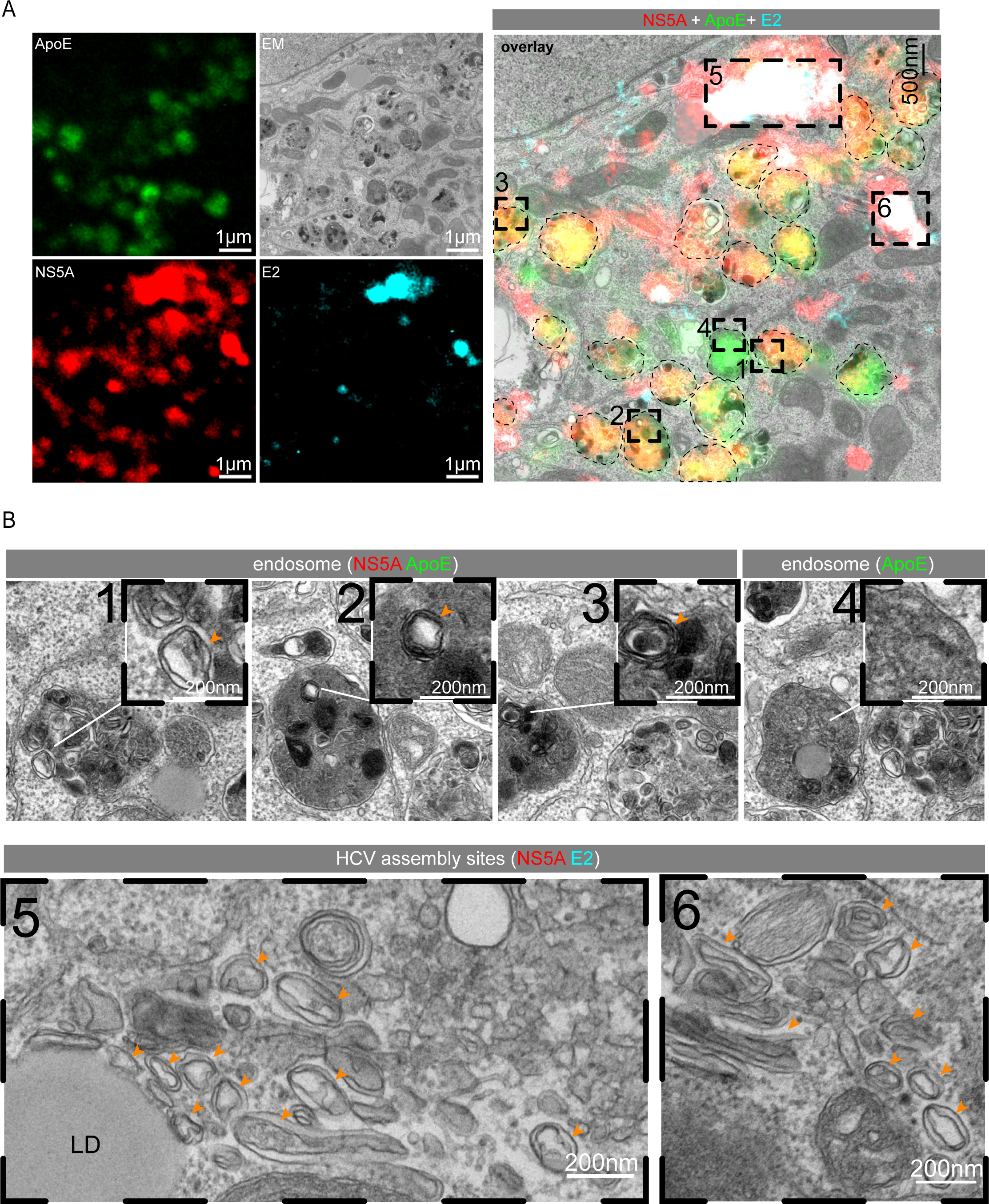
Detection of HCV-produced intraluminal vesicles in NS5A-ApoE double-positive endosomes. (A) Huh7-Lunet/ApoE^mT2^ cells expressing HCV Core-NS2/E2^eYFP^ and containing the subgenomic replicon sgJFH1/NS5A^mCherry^ (Fig. 2A) were investigated with the CLEM method at 48 h p.e. Lipid droplets stained with lipidTox were used as fiducial markers to correlate light and electron micrographs. Dashed squares in the overlay image (right panel) refer to NS5A-ApoE double-positive structures. The left panels show single-channel light or EM micrographs of the enlarged overlay image on the right. For ease of visualization, endosome peripheries are marked with dashed lines. (B) Magnified views of regions indicated in the dashed squared areas in the overlay image in (A). Cropped areas 1, 2, 3: ApoE-NS5A double-positive ILVs in endosomes; crop 4: an ApoE-positive, NS5A-negative endosome; cropped areas 5 and 6: NS5A-E2 double-positive areas containing numerous DMVs. Orange arrowheads point to ILVs in crops 1, 2 and 3; and DMVs in crops 5 and 6. LD, lipid droplet.

### Co-secretion and co-transmission of ApoE-positive lipoproteins with endosome-derived extracellular vesicles containing HCV NS5A and viral RNA

ApoE associates with NS5A in regions of endosomes containing HCV-produced intraluminal double or multi-membrane vesicles (Figs 5 and 6). Moreover, HCV suppresses the fusion of late endosomes with lysosomes (67). Therefore, we hypothesized that secreted ApoE-LPs might associate with HCV-produced EVs containing NS5A and viral HCV RNA. To address this assumption, we employed a subgenomic HCV replicon that supports viral RNA transmission via endosome-derived EVs, albeit with a rather low efficiency (33–36). In the first set of experiments, we determined whether ApoE associates with NS5A and viral RNA released from cells containing a stable subgenomic HCV replicon or parental control cells by using ApoE-specific pull-down. Captured complexes were analyzed by HCV-specific RT-qPCR. As shown in Fig 7A, we detected HCV RNA in immuno-captured ApoE-containing complexes isolated from supernatants of replicon-containing cells. Samples captured with control antibodies or from mock-transfected cells were at the background level arguing for the release of EVs containing viral RNA from replicon cells.

**Fig 7.**
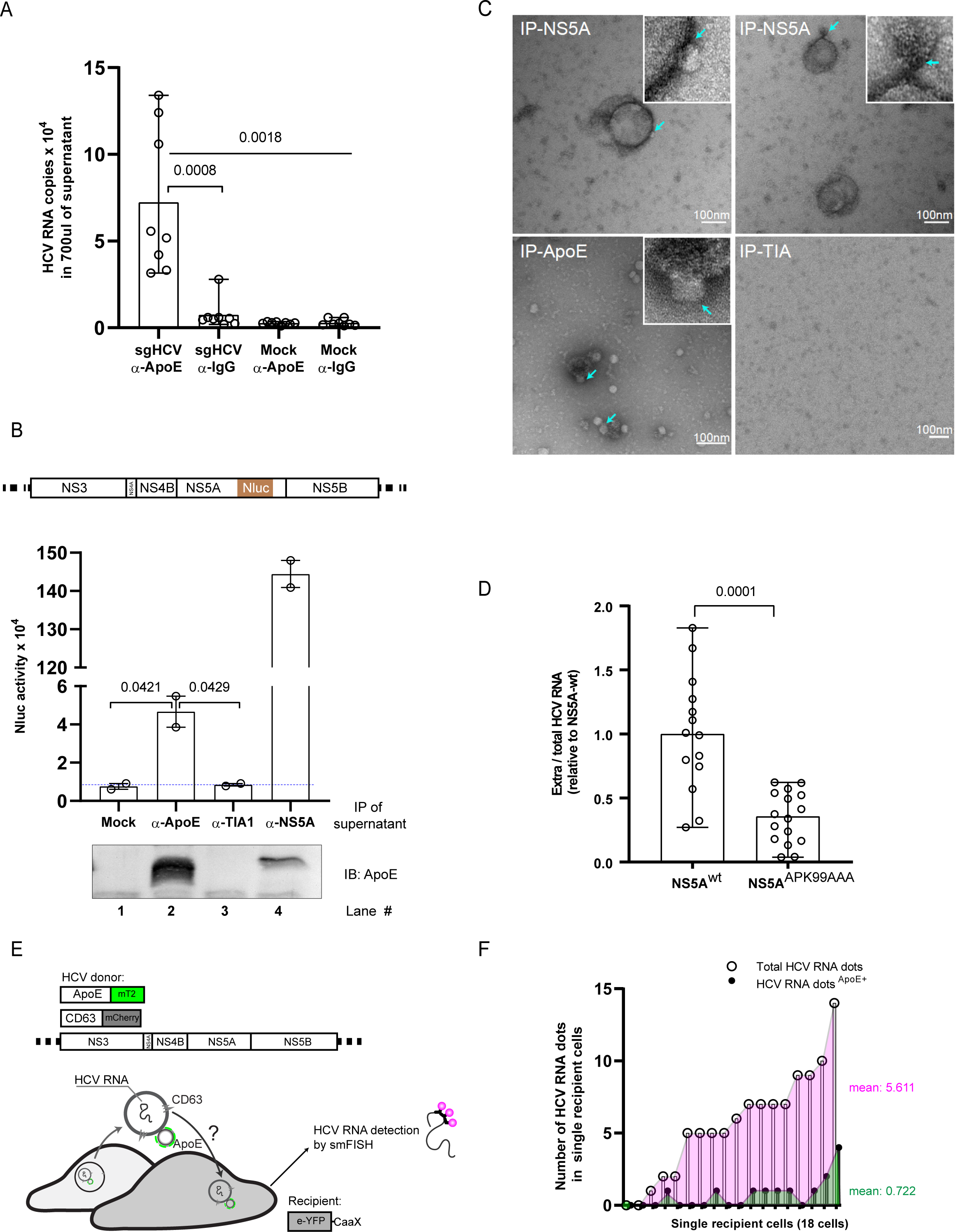
Co-secretion and co-transmission of ApoE-LPs with endosome-derived EVs containing HCV NS5A and RNA. (A) Virion-free release of HCV RNA in association with ApoE. Huh7 cells harboring a subgenomic HCV replicon and control cells were cultured in a medium containing 1% FCS for 6 h. Culture supernatants were subjected to immunoprecipitation using ApoE-specific or IgG control antibodies. Immuno-complexes were analyzed by HCV-specific RT-qPCR. Data are means (range) from 2 independent experiments. Single dots represent technical replicates from the two experiments. P-value was determined using one-way ANOVA and unpaired Student’s *t*-tests. (B-C) Association of secreted ApoE with NS5A-containing EVs. (B, top panel) Schematic of the Nanoluciferase (Nluc)-tagged NS5A subgenomic replicon construct. (B, middle and bottom panel) Huh7-Lunet cells were electroporated with subgenomic replicon RNA encoding the NLuc-tagged NS5A and 72 h p.e, culture supernatant was subjected to immunoprecipitation using ApoE-, or NS5A-, or control TIA1-specific antibodies. NS5A contained in captured immuno-complexes was quantified by measuring NLuc activity (middle panel). ApoE contained in captured complexes was analyzed by Western blot (bottom panel). Data are means (range) of two independent experiments. P-value was determined using unpaired Student’s *t*-test. (C) Captured complexes from (B) were visualized by negative staining and analyzed by TEM. Turquoise arrows point to LP-like particles (∼20 nm) attached to EVs that were captured with antibodies specified on the top of each panel. (D) Reduced virion-free secretion of HCV RNA with the ApoE-binding deficient NS5A^APK99AAA^ mutant. Total RNA contained in Huh7-Lunet cells with stable wildtype or mutant subgenomic replicon was extracted and HCV RNA was quantified by RT-qPCR. In addition, total RNA in culture supernatants was isolated and HCV RNA contained therein was quantified by RT-qPCR. Ratios of secreted to total HCV RNA are shown. Data are medians (range) from three independent experiments. Single dots represent technical replicates of the 3 biological experiments. P-value was determined using unpaired Student’s *t*-test. (E-F) Uptake of ApoE-associated, virion-free released HCV RNA by HCV-negative bystander cells. (E) Experimental approach. Huh7-Lunet cells expressing tagged ApoE and CD63 and containing a subgenomic replicon (constructs on the top) served as donor cells. Huh7-Lunet-derived recipient cells expressed eYFP, fused to the farnesylation signal from human HRAS protein (CaaX) to visualize cellular membranes. Donor and recipient cells were co-cultured for 24 h, fixed, and HCV RNA in recipient cells was detected by using smFISH with Hulu probes. (F) Number of total and ApoE-positive HCV RNA dots in each analyzed cell (n=18) is shown.

To verify the presence of NS5A in the ApoE-captured complexes, we transfected Huh7-Lunet cells with a subgenomic replicon RNA encoding Nanoluciferase (Nluc)-tagged NS5A to allow its sensitive detection in cell culture supernatants (Fig 7B, upper). In agreement with a previous report (60), we observed time-dependent secretion of NS5A^Nluc^ into the cell culture supernatant (Fig 4F). Importantly, Nluc activity was clearly detected upon ApoE-specific immunocapture indicating a direct or indirect association between NS5A and ApoE (Fig 7B, lane 2). The specificity of the pull-down was confirmed by using mock cells or an unrelated antibody (Fig 7B, lane 1 and 3, respectively). Surprisingly, the highest Nluc activity was detected in NS5A-captured immunocomplexes, arguing that NS5A is well-accessible on the outside of EVs (Fig 7B, lane 4). Negative-staining of immunocaptured samples confirmed that NS5A- and ApoE-associated structures correspond, at least in part, to EVs that were frequently associated with LP-like structures (Fig 7C, arrows).

Next, we examined the possible relevance of ApoE-NS5A interaction for the secretion of EVs containing HCV RNA. To this end, we used an NS5A mutant (APK99AAA) reported to have a defect in interaction with ApoE (Fig S4G) (57). Of note, Huh7-Lunet cells containing a stable subgenomic replicon encoding mutant NS5A^APK99AAA^ released much lower amounts of HCV-RNA than the wildtype replicon (Fig 7D). These results suggest that ApoE - NS5A interaction is required for the efficient release of EVs containing viral RNA, providing an explanation for the association of ApoE with NS5A-positive EVs released from HCV-replicating cells.

With the aim to visualize intracellular HCV RNA and its association with ApoE, we employed single molecule Fluorescence In Situ Hybridization (smFISH). Used probes were conjugated to Alexa Fluor 647 and enabled visualization of single HCV RNA molecules without signal amplification (Fig S5A). In spite of some nuclear background staining, cytoplasmic staining of HCV RNAs was specific as we detected numerous cytoplasmic foci of viral RNA in replicon cells, but not in the control cells (Fig S5B). To determine if ApoE associates with HCV RNA-containing EVs that might be transferred to neighboring (bystander) cells, we established Huh7-Lunet/ApoE^mT2^ cells containing a stable subgenomic HCV replicon and expressing CD63^mCherry^ (Fig 7E). These cells were used for co-culture experiments and served as donors. As recipient cells, we used HCV-negative Huh7-Lunet cells expressing the CaaX^eYFP^ membrane sensor (see Fig 3C). HCV RNAs were found to partially colocalize with ApoE-CD63 double-positive puncta in donor cells (Fig S5C, area 1). Remarkably, we could detect distinct foci of HCV RNA in single recipient cells, around 13% of them being ApoE-CD63 double-positive (example image in Fig S5C, area 2; quantification in Fig 7F). These data suggest that HCV might hijack the late endosomal trafficking and egress of ApoE-LPs to transmit NS5A and viral RNA via endosome-derived EVs.

## Discussion

In this study, we developed two tags for ApoE labeling that do not impact its function while allowing the tracking of hepatocyte-made ApoE by live-cell imaging and various other imaging modalities. Obtained results suggest that hepatic ApoE-LPs follow the trafficking pathway of CD63-positive late endosomes. This pathway appears to be hijacked by HCV using the multi-functional protein NS5A that binds to ApoE to release EVs containing viral RNA. Our observations suggest that late endosomes in hepatocytes might be a central site for the storage and secretion of ApoE-LPs. Since biosynthesis and secretion of ApoE-LPs such as VLDL depend largely on the availability of dietary fat, and have to respond rapidly to elevated plasma insulin levels by retaining hepatic lipids (68, 69), a lipid reservoir like late endosomes would allow rapid response to fluctuating food and insulin levels.

Several viruses exploit ApoE for their replication cycles. Two prominent examples are HBV and HCV that both associate with ApoE-containing lipoprotein particles (17, 32, 70). In the case of HCV, ApoE interacts with NS5A and the envelope glycoproteins and these interactions are critical for HCV particle assembly and maturation (32, 57, 58). Here, we provide evidence that ApoE-NS5A interaction is additionally required for the secretion of HCV-induced EVs containing viral RNA. Release of HCV NS5A and virion-free RNA has been suggested in several independent studies (33-36, 59, 60, 71), but the role of ApoE in this process has not been studied. Our results suggest that ApoE is a critical component for the release of EVs from HCV-replicating cells and these vesicles can be transmitted to naïve bystander cells, consistent with the virion-free transfer of intact HCV genomes from cell to cell (33–36).

Our results address another long-standing conundrum in HCV biology, i.e. the tight association between ApoE and NS5A (56–58). Although both proteins localize to opposing sites of the ER membrane (28, 72), we can efficiently capture EVs from HCV-replicating cells by NS5A pull-down, indicating that NS5A resides on the surface of EVs where it can interact with ApoE. How NS5A might end up on the surface of these vesicles is not known. For the poliovirus it has been shown that the viral replicase complex resides on the surface of the replication vesicles, which are double-membrane vesicles like for HCV, and a similar topology might apply to NS5A (73–75). Regarding the functional relevance, we note that the ApoE-NS5A interaction is not required for HCV virion assembly, at least in the subgenomic replicon model, but appears to boost the release of viral RNA from infected cells, e.g. to avoid recognition by innate RNA sensors such as TLR3 (59).

EVs are phospholipid bilayer-enclosed structures released from cells and containing various signaling molecules (76–79). They are considered as a “language” exploited by cells and viruses for intercellular communication (80–83). Several lines of evidence argue for interaction between LPs and EVs. First, various procedures of EV isolation and purification, including size and density fractionation as well as enrichment of CD63-positive EVs do not allow complete separation of LPs and EVs (52–54, 84). Second, LPs were found to attach *in vitro* to purified EVs or even fuse to crude EVs in blood plasma (20, 21, 85, 86). Third, pigment cell-derived ApoE associates with endosome-derived ILVs and plays an important role in the sorting of a distinct cargo to ILVs and its release via exosomes (30). Although these studies suggest an association of LPs with ILVs/EVs, to the best of our knowledge, the association between liver-generated LPs and endosome-derived EVs is not well documented and their possible intercellular co-transmission has been unknown.

Our data suggest that in naïve and HCV-infected hepatocytes, ApoE-LPs and endosome-derived CD63-positive ILVs/EVs not only share a common intracellular late endosomal trafficking route, but also are partially co-secreted. These particle complexes forming intracellularly co-enter target cells, arguing for a stable interaction between ApoE-LPs and CD63-positive ILVs/EVs. This would explain the difficulty to separate LPs from EVs (52–54, 84), which poses a major challenge to assign distinct functions to each of these vesicle species individually (87). The mechanism underlying this interaction is unknown, but might be mediated by associations between ApoE on LPs and scavenger receptor class B type 1 (SR-BI) or heparan sulfate decorating the surface of ILVs/EVs (86, 88). These interactions could also modulate lipid transfer from LPs to ILVs/EVs (86). Moreover, since hepatic ApoE-LPs are secreted into the blood stream, they might alter the systemic spread of EVs into different distant tissues and organs, thus manipulating various biological responses depending on EV content. For instance, the amount of liver-generated plasma ApoE was found to be associated with unfavorable alterations in neurodegenerative diseases including synaptic integrity (89). The underlying mechanism has not been determined but might be due to the direct contribution of ApoE to lipid metabolism or ApoE-facilitated blood-brain barrier passage of EVs (90–93). Another example is COVID-19 where plasma-derived EVs isolated from COVID-19 patients alter multiple signaling pathways (94), which might contribute to the broad spectrum of clinical symptoms (95). Importantly, COVID-19 derived EVs preparations contain multiple apolipoproteins including ApoE, ApoB, ApoA2, ApoD, and ApoH (94).

Our study has some limitations. It is primarily based on the use of human hepatoma cells that are highly permissive to HCV and easy to manipulate. However, because LPs and ILV/EV profiles in vivo are somewhat different, future studies require more physiologically relevant systems, which are however, not permissive to HCV and difficult to manipulate. In addition, although the HCV subgenomic replicon model allows excluding the transmission of HCV RNA via virions, HCV-produced ILVs/EVs might also contain viral structural proteins including the envelope glycoproteins E1 and E2, potentially assisting in the spread of these vesicles (96). Finally, the physiological consequences of co-spread of hepatic LPs with -EVs in general and in the context of HCV infection, the latter possibly allowing HCV RNA spread independent of virus particles (33–36) remain to be determined but they are beyond the scope of the present study.

In conclusion, our study provides insights into the endosomal egress and transmission of hepatocyte-derived ApoE-containing LPs and the strategy how HCV exploits this pathway. Given the more general role of EV-mediated cell-to-cell communication, the association of ApoE-LPs with EVs reported here provides new starting points for research into the pathophysiology of ApoE-related metabolic and infection-related disorders.

## Materials and Methods

### Materials

Reagents and resources used in this study are provided in Table S1.

## Methods

### Cell lines and culture conditions

All cells used in this study were cultured in Dulbecco’s modified Eagle medium (DMEM, Thermo Fisher Scientific), supplemented with 2 mM L-glutamine, nonessential amino acids, 100 U/ml of penicillin, 100 µg/ml of streptomycin, 10% fetal calf serum (DMEMcplt) and given concentrations of antibiotics to select for stable expression of genes of interest. Huh7-Lunet/CD81H cells (750 µg/ml G418) derived from the Huh7 subclone Huh7-Lunet (97) and expressing high levels of the HCV entry receptor CD81, and Huh7-Lunet/CD81H/ApoE-KD cells (5 µg/ml puromycin) with a stable knockdown of ApoE have been described earlier (32, 38). For reasons of simplicity, in this study Huh7-Lunet/CD81H cells are designated Huh7-Lunet cells. HEK293T-miR122 cells (2 µg/ml puromycin), kindly provided by Thomas Pietschmann, have been reported elsewhere (40). Huh7.5 and HEK293T cells have been described elsewhere (98, 99). HEK293T-miR122, Hela Kyoto, and Huh7-Lunet/ApoE-KD cells were used to generate ApoE^mT2^ expressing cells by lentiviral transduction and stable selection with 10 μg/ml blasticidin. For the production of HCV-like transcomplemented particles (HCV_TCP_), Huh7-Lunet/ApoE-KD/ApoE^mT2^ cells (designated Huh7-Lunet/ApoE^mT2^ in this study for reasons of simplicity) were transduced with lentiviruses encoding the HCV structural proteins (C-E1-E2^eYFP^-p7-NS2 or C-E1-E2^eYFP^-p7), selected with 500 μg/ml Zeocin and maintained in 50 ug/ml Zeocin-containing DMEMcplt. To obtain cells with stably replicating subgenomic replicon of the HCV strain JFH1 and used for the coculture experiment, Huh7-Lunet/ApoE^mT2^/CD63^mCherry^ cells were electroporated with *in vitro* transcripts of the construct sgHyg/JFH1. To monitor HCV RNA secretion in the context of an ApoE-binding defective NS5A mutant or wildtype NS5A, Huh7-Lunet cells were electroporated with *in vitro* transcripts of the construct sgHyg/JFH1/NS5A^APK99AAA^ or sgHyg/JFH1, respectively. Stable cells were selected in a medium containing 400 μg/ml hygromycin and maintained in 150 µg/ml hygromycin-containing DMEMcplt. FCS devoid of extracellular vesicles (EVs) was prepared as previously described (59). The full names of constructs used in this study are given in the Supporting Table 1.

### Antibodies and immunofluorescence reagents

All antibodies and immunofluorescence reagents used in this study are listed in S1 Table.

### DNA plasmid constructs

The lentiviral construct pWPI_ApoE encoding human ApoE3 was described previously (100). To generate pWPI_ApoE^FP^ and pWPI_ApoE^SNAPf^ constructs, the FP- and the SNAPf-coding sequences were amplified by PCR using the corresponding plasmids as templates (see Supporting Table 1) and inserted at the 3’ end of the ApoE-coding sequence via the linker sequence SGGRGG. Construct pWPI_CD63^mCherry^ encodes a fusion protein of human CD63 and C-terminal mCherry. To generate the construct pWPI_eYFP-CaaX, the eYFP-coding sequence was extended at the 3’ end by the CaaX coding sequence derived from the human HRAS protein and inserted into the lentiviral vector pWPI. To generate pWPI_CD63_M153R_pHluorin, the CD63-pHluorin coding sequence contained in plasmid pCMV-Sport6-CD63-pHluorin (51) was amplified by PCR and inserted into the lentiviral vector pWPI. To stabilize pHluorin and increase signal intensity, we inserted the M153R mutation (50) by using PCR and primers carrying the desired nucleotide substitutions.

The full-length HCV constructs Jc1 and JcR2A have been described elsewhere (42, 101). The lentiviral constructs encoding the HCV structural proteins Core-NS2/E2^eYFP^ or Core-p7/E2^eYFP^ were created by replacing the eGFP-coding sequence reported previously (62) by the eYFP-coding sequence. Plasmid pFK_I389neoNS3-3′_dg_JFH1_NS5A-aa2359_mCherry_NS3-K1402Q (designated sgNeo/JFH1/NS5A^mcherry^ in this study) has been reported earlier (102). To generate the subgenomic replicon encoding a CLIPf-tagged NS5A and the neomycin resistance gene (construct sgNeo/JFH1/NS5A^CLIPf^), the mCherry-coding sequence in construct sgNeo/JFH1/NS5A^mcherry^ was replaced by the CLIPf-coding sequence. To allow selection with hygromycin, the neomycin resistance gene was replaced by the hygromycin resistance gene. To generate the subgenomic replicon construct encoding a NanoLuciferase-tagged NS5A (sgHyg/JFH1/NS5A^Nluc^), the mCherry-coding sequence of construct sgHyg/JFH1/NS5A^mCherry^ was replaced by the NanoLuciferase-coding sequence (103). The mutations in NS5A interfering with ApoE interaction (APK99AAA) (57, 104) were inserted into the replicon construct sgHyg/JFH1 by using PCR-based mutagenesis.

To generate plasmids encoding myc-tagged NS5A wildtype and the APK99AAA mutant corresponding plasmids were used as template for PCR using primers encoding the myc-tag sequence and NS5A sequences were inserted into the pCDNA3^+^ vector. Other plasmids used in this study are listed in the Supporting Table 1.

### Preparation of *in vitro* transcripts and electroporation of HCV RNA

HCV RNA preparations generated by *in vitro* transcription and transfection of cells by electroporation have been described elsewhere (105). In brief, plasmids containing HCV JFH1 genomes were linearized using the restriction enzyme MluI-HF (NEB) and purified using the NucleoSpin Extract II Kit (Macherey-Nagel). RNA transcripts were synthesized via *in vitro* transcription using T7 RNA polymerase in 100 µl-reaction mixtures [80 mM HEPES (pH 7.5), 12 mM MgCl_2_, 2 mM spermidine, 40 mM dithiothreitol, 3.125 mM of each rNTP, 1 U/µl RNasin (Promega), 0.6 U/µl T7 RNA polymerase, and the respective linearized DNA template]. After 4 h at 37°C, the DNA template was degraded by 45 min treatment with 2 U of RNase-free DNase (Promega) per 1 µg DNA at 37°C. RNA was purified by acidic phenol-chloroform extraction, precipitated with isopropanol, and dissolved in RNase-free water. The integrity and concentration of RNA were evaluated using agarose gel electrophoresis and spectrophotometry.

For electroporation, confluent cell monolayers were trypsinized and resuspended in Cytomix [120 mM KCl, 0.15 mM CaCl_2_, 10 mM potassium phosphate buffer, 25 mM HEPES (pH 7.6), 2 mM EGTA, and 5 mM MgCl_2_] (106) containing 2 mM ATP and 5 mM glutathione (1-2×10^7^ cells/ml). *In vitro* transcripts (5 µg) were mixed with 200 µl of the cell suspension and electroporation was performed at 975 µF and 166 V using the Gene Pulser system (Bio-Rad) and a cuvette with a gap width of 2 mm (Bio-Rad). Alternatively, 10 µg *in vitro* transcripts were mixed with 400 µl of the cell suspension and electroporation was performed at 975 µF and 270 V using a cuvette with a gap width of 4 mm. After electroporation, cells were immediately transferred to DMEMcplt and seeded into the desired cell culture dishes.

### Western blot analysis

Cell extracts were prepared using 2x sample buffer [120 mM Tris-HCl (pH 6.8), 60 mM SDS, 100 mM DTT, 1.75% glycerol, 0.1% bromophenol blue] supplemented with 5 mM MgCl_2_ and 5 U/ml benzonase. Samples were denatured by heating to 95°C for 5 min. Proteins were separated by SDS-PAGE and transferred to a polyvinylidene difluoride (PVDF) membrane that was blocked by incubation in 5% skim milk-containing PBS-0.05% Tween 20, pH 7.4 (PBST) for 1 h at room temperature (RT). The membrane was incubated with a primary antibody in 1% skim milk-containing PBST for either 1 h at RT or overnight at 4°C and subsequently incubated with a secondary antibody conjugated with horseradish peroxidase (HRP) for 1 h at RT. Bound secondary antibodies were detected using the Western Lightning Plus-ECL reagent (PerkinElmer) and signals were visualized by using the Intas ChemoCam Imager 3.2 (Intas).

### Quantitative detection of HCV RNA by RT-qPCR

Total RNA contained in cell lysates or cell culture supernatant was extracted using the NucleoSpin RNA extraction kit (Macherey-Nagel) according to the instruction of the manufacturer. HCV RNA copy numbers in extracted samples were determined with HCV-specific primers and a probe by using the Quanta BioSciences qScript XLT One-Step RT-qPCR KIT (Quanta Biosciences, Gaithersburg, MD) as described elsewhere (59). Serially diluted HCV *in vitro* transcripts were included in parallel to calculate HCV RNA copy numbers contained in analyzed samples.

### Quantification of HCV Core protein

HCV core protein amount was quantified using a commercial Chemiluminescent Microparticle Immunoassay (CMIA) (6L47, ARCHITECT HCV Ag Reagent Kit, Abbott Diagnostics) as reported earlier (62).

### Production of lentiviruses

Lentiviruses encoding genes of interest were produced as described recently (107). In brief, HEK-293T cells were co-transfected with the human immunodeficiency virus-Gag packaging plasmid pCMV-dR8.91, the vesicular stomatitis virus-G encoding plasmid pMD2.G, and the pWPI construct containing the gene of interest by using polyethylenimine (Polysciences Inc.). Lentivirus-containing supernatants were harvested at about 48 h post-transfection and filtered through a 0.45 μm pore-size filter (MF-Millipore).

### Live-cell time-lapse confocal microscopy

Cells were seeded onto either 4-compartment (CELLview, Greiner BIO-ONE) or 1-compartment (MatTek Corporation) 35 mm-diameter glass-bottom imaging dishes. Prior to imaging, cells were washed twice and cultured in phenol red-free DMEMcplt. Live-cell time-lapse confocal microscopy was performed in a humidified incubation chamber at 37°C and 5% CO_2_ using a PerkinElmer UltraVIEW Vox Spinning Disc microscope equipped with Yokogawa CSU-X1 spinning disk head, Nikon TiE microscope body, a Hamamatsu C9100-23B EM-CCD camera, an automated stage and the Perfect Focus System (PFS). An Apo TIRF 60x/1.49 N.A. oil immersion objective was used. Multichannel images were acquired sequentially using solid state lasers with excitation at 445 nm for mTurquoise2, 488 nm for pHluorin, 514 nm for eYFP, 561 nm for CLIP^ATTO590^, 640 nm for SNAP^SIR647^, and matching emission filters. For imaging of pHluorin-tagged CD63 expressing cells, the medium was supplemented with 25 mM Hepes (pH 7.4) to stabilize neutral pH. The imaging time interval of each experiment is specified in the figure legends.

### Immunofluorescence staining and confocal microscopy

Immunofluorescence (IF) staining was performed as previously described (32). Briefly, cells seeded onto coverslips were fixed with 4 % paraformaldehyde (PFA) in PBS for 10 min at RT and permeabilized with 0.1% Triton X-100 in PBS for 10 min at RT. After blocking with 3% (w/v) bovine serum albumin (BSA) in PBS for 20 min at RT, cells were incubated with a diluted primary antibody in 1% BSA/PBS for 1 h at RT or overnight at 4°C. Cells were further incubated with a diluted secondary antibody conjugated with an Alexa fluorophore (1:1000) in 1% BSA/PBS (Molecular Probes) for 1 h in a dark condition at RT. If required, cell nuclei were counterstained with DAPI (1:3000) (Molecular Probes). In-between each step, cells were washed at least 3 times with 1x PBS. Unless otherwise stated, coverslips were mounted with Fluoromount-G mounting medium (Electron Microscopy Sciences, Ft. Washington, USA) overnight at 4°C. For selective permeabilization assay, cells were permeabilized in 5 µg/ml digitonin dissolved in PBS for 15 min at 4°C. IF images were generated with a spinning disc confocal microscope (PerkinElmer).

### Super-resolution microscopy

Huh7-Lunet/ApoE^SNAPf^ cells stably expressing ApoE^SNAPf^ were electroporated with *in vitro* transcripts of HCV sgNeo/JFH1/NS5A^CLIPf^ and grown on high precision glass coverslips (Deckglaeser, Marienfeld). At 48 h post-electroporation, cells were sequentially incubated with CLIP^ATTO590^ (1:2500) and 5 µM SNAP^SiR647^ (NEB) in DMEMcplt for 1 h. Cells were washed intensively at least 3 times with DMEMcplt and cultured for 15 min. Thereafter, cells were washed 3 times with PBS, fixed with 4% PFA in PBS for 10 min at RT, and subjected to immunofluorescence staining using anti-CD63 antibody conjugated to Alexa Fluor 488 (Santa Cruz). Cells were mounted with ProLong Gold Antifade Mountant (ThermoFisher Scientific) by overnight incubation at RT. STED imaging was conducted using an Expert Line STED system (Abberior Instruments GmbH, Göttingen, Germany) equipped with an SLM based easy3D module, an Olympus IX83 microscope body, solid state pulsed lasers (488 nm, 590 nm, and 640 nm), and the 775 nm STED laser. The 100x oil immersion objective (NA, 1.4; Olympus UPlanSApo) was used. Initially, confocal images were captured in the line sequential mode using the following excitation lasers: 488 nm for AF488, 590 nm for ATTO590, 640 nm for SIR647, and the corresponding 525/50, 615/20, and 685/70 emission filters. These filters are placed in front of avalanche photodiodes for detection. Small regions of interest were selected and subjected to STED imaging. STED images in selected areas were captured sequentially using the 590 nm and 640 nm excitation laser lines in the line sequential mode with corresponding 615/20 and 685/70 emission filters, followed by the depletion using the 775 nm STED laser. STED images were deconvoluted using the Huygens Deconvolution software (Scientific Volume Imaging) using Classic Maximum Likelihood Estimation (CMLE) algorithm and Deconvolution Express mode with “Conservative” settings.

### HCV RNA detection by single-molecule fluorescence in situ hybridization

Intracellular HCV RNAs were visualized by smFISH using Hulu probes (PixelBiotech, Germany) according to the manufacturer’s instruction with slight modifications. In brief, cells grown on glass coverslips were fixed with 4% paraformaldehyde (PFA) in PBS for 30 min at RT. Cells were then treated with 150 mM glycine in PBS to quench residual PFA, permeabilized by treatment with 0.1% Triton X-100 in PBS for 10 min, and incubated with proteinase K (1:4000) (ViewRNA ISH Kit, ThermoFisher Scientific) in PBS for 5 min. HCV RNAs were hybridized to Hulu probes targeting the positive strand in the NS3 coding region (nucleotides 3733-4889 of the JFH-1 genome; GenBank accession number AB047639). Hybridization was done in HuluHyb solution (2xSSC, 2 M Urea, 10% dextran sulfate, 5x Denhardt’s solution) using a humidified chamber at 30°C overnight. Cells were washed extensively with HuluWash and coverslips were mounted on glass slides with Prolong Gold Antifade Mountant (ThermoFisher Scientific) by overnight incubation at RT.

### Immunoprecipitation

HEK293T-miR122 cells were co-transfected with HA-tagged ApoE construct, or an empty vector, or pCDNA3+ myc-tagged NS5A^wt^, or myc-tagged NS5A^APK99AAA^, respectively, using the TransIT-LT1 Transfection Reagent (Mirus Bio). After 30 h, cells were lysed by 10 min incubation in lysis buffer [50 mM Tris-HCl, pH 7.5, 150 mM NaCl, 1% Triton X-100, 1 mM EDTA, 10% glycerol, 1x protease inhibitor cocktail (Roche)] on ice. Cell lysates were centrifuged at 15,000 x *g* for 15 min at 4°C. Cleared supernatants were incubated with protein G-magnetic bead slurry (Dynabeads, ThermoFisher Scientific) for 30 min at 4°C to remove proteins binding to the resin. Beads were removed by pelleting with a magnetic stand and supernatants were incubated with rabbit anti c-myc antibody at 4°C overnight. Protein complexes were captured by adding protein G bead slurry and 1 h incubation of samples under continuous rotation at 4°C. Beads were washed 5 times with lysis buffer lacking glycerol, captured protein complexes were eluted with 2x sample buffer and denatured for 5 min at 95°C. Proteins were analyzed by Western blot using mouse anti-HA antibody.

### Iodixanol density gradient centrifugation

Cells were washed and cultured for 5 h in 1% FCS-containing DMEM. Thereafter, cell culture supernatant was filtered through a 0.45 μm pore-size filter (MF-Millipore), loaded on top of a PBS-based 10-50% iodixanol gradient (Sigma Aldrich), and subjected to isopycnic centrifugation for 18 h at 34,000 rpm (∼120,000 x *g*) at 4°C using an SW60 rotor (Beckman Coulter, Inc.). Eleven fractions were collected from top to bottom and analyzed by density measurement using a refractometer (Kruess, AGS Scientific) and Western blot.

### Luciferase reporter assay

HCV RNA replication kinetics were determined by using the HCV JcR2A reporter construct. Briefly, cells were collected at 4, 24, 48 and 72 h post-electroporation and lysed in luciferase lysis buffer (1% Triton X-100, 10% glycerol, 25 mM glycylglycine, 15 mM MgSO_4_, 4 mM EGTA, and 1 mM DTT) for 15 min at RT. Cell lysates were transferred to 96-well plates and coelenterazine-containing luciferase assay buffer (25 mM glycylglycine, 15 mM MgSO_4_, 4 mM EGTA, 1 mM DTT, and 15 mM K_3_PO4, pH 7.8) was injected. Renilla luciferase activities were measured using a Mithras LB 940 plate luminometer (Berthold Technologies, Freiburg, Germany). Obtained values were normalized to the 4 h value of each transfection to correct for transfection efficiency. To measure the transmission of HCV, culture supernatants were used to inoculate naïve Huh7.5 cells, and after 72 h, cells were lysed and subjected to luciferase assay. NanoLuciferase (Nluc) activity was measured using the Nano-Glo Luciferase Assay System (Promega) according to the instruction of the manufacturer with slight modifications. In brief, 50 µl of samples were mixed with 50 µl NLuc substrate (1:1000) in the assay buffer and NLuc activities were measured using a Mithras LB 940 plate luminometer (Berthold Technologies, Freiburg, Germany).

### Immunocapture of extracellular ApoE-associated structures

Supernatants of cells cultured in EV-free DMEM were collected, filtered through a 0.45 μm pore-size filter (MF-Millipore), and incubated with an anti-ApoE antibody for 3 h at 4°C. ApoE-associated structures were captured using protein G-magnetic beads (Dynabeads, ThermoFisher Scientific) and overnight incubation at 4°C with continuous rotation. After 5 times washing with ice-cold PBS, protein complexes were eluted by 10 min incubation with 0.1 M glycine, pH 2.5 at RT, and samples were neutralized by adding 1 M Tris, pH 7.5.

### Transmission electron microscopy and correlative light and electron microscopy (CLEM)

Sample preparation, data acquisition, and data processing were conducted as described earlier (62) with slight modifications. For CLEM, cells were fixed for 30 min at RT with a fixative containing 0.2% glutaraldehyde (GA) and 4% PFA and then washed 3 times with PBS to remove the fixative. The coordinates of cells-of-interest on the gridded MatTek dish were captured with the 20x objective using transmitted light with differential interference contrast (DIC). Cells were then subjected to immunofluorescence imaging using an oil immersion 60x objective, covering the ∼2.8 µm cell thickness with 0.2 µm spacing between optical planes before and after the addition of LipidTox™ Deep Red Neutral Lipid Stain (Invitrogen). Cells were further postfixed in 2.5% GA in CaCo buffer with supplemented ions [2.5% GA, 2% sucrose, 50 mM sodium cacodylate (CaCo), 50 mM KCl, 2.6 mM MgCl_2_, and 2.6 mM CaCl_2_] for 30 min or overnight at 4°C. After 3 washes with 50 mM CaCo buffer, cells were incubated with 2% osmium tetroxide in 50 mM CaCo for 40 min on ice, washed 3 times with milli-Q water, and incubated with 0.5% uranyl acetate in water at 4°C. Samples were washed again with water prior to the sequential dehydration of cells using a graded ethanol series from 50% to 100% at RT. Samples were embedded in Epon 812 (Carl Roth) and incubated for at least 2 days at 60°C to allow polymerization of the resin. Epon was detached from the glass coverslips by dipping it several times into liquid nitrogen followed by hot water. Cells of interest were identified by the negative imprint of the gridded coverslips and cut into 70 nm ultrathin sections using an ultramicrotome (Leica EM UC6, Leica Microsystems). Sections were collected on pioloform coated copper palladium slot grids (Science Services, GMBH) and counterstained sequentially with 3% uranyl acetate in water for 5 min and lead citrate (Reynold’s) for 5 min. Images were acquired by using the Jeol JEM-1400 (Jeol Ltd., Tokyo, Japan) transmission electron microscope (TEM) equipped with a 4k pixel digital camera (TemCam F416; TVIPS, Gauting, Germany) and the EM-Menu or Serial EM software (108). Lipid droplets were used as fiducial markers to correlate the EM with the light micrographs using the Landmark Correspondences plugin in the Fiji software package (109). To visualize ApoE-containing structures enriched by immunocapture, samples were added onto freshly glow-discharged carbon- and pioloform-coated 300-mesh copper grids (Science Services GmbH, Munich, Germany) and subjected to negative staining using 3% uranyl acetate for 5 min at RT.

### Immunogold labeling

For immunogold labeling of ApoE-associated structures, all incubation and washing steps were conducted by floating the grids on top of drops at RT. In-between each step, samples were washed at least 5 times for 2 min with PBS. The basic protocol employed has been reported elsewhere (110) and only slight modifications were made. In brief, samples absorbed onto copper grids were blocked with the blocking solution [0.8% BSA (Roth, Karlsruhe, Germany), 0.1% fish skin gelatin (Sigma-Aldrich), 50 mM glycine in PBS]. For ApoE and CD63 labeling, grids were incubated with goat anti-ApoE antibody (1:100) and mouse anti-CD63 antibody (1:100) in blocking solution, respectively, for 30 min at RT. Grids were further incubated with rabbit anti-goat- or anti-mouse-bridging antibody (1:150) in the blocking solution for 20 min. Bound antibodies were detected with protein A conjugated to 10-nm gold particles diluted 1:50 in blocking buffer for 30 min. Grids were fixed with 1% glutaraldehyde in PBS for 5 min, washed 7 times with H_2_O, briefly rinsed with 3% uranyl acetate, and negatively stained again with 3% uranyl acetate for at least 5 min.

### Automated particle tracking in fluorescence microscopy images

Particle tracking in fluorescence microscopy images was performed by using a probabilistic particle tracking approach that is based on Bayesian filtering and multi-sensor data fusion (111). This approach combines Kalman filtering and particle filtering and integrates multiple measurements by separate sensor models as well as sequential multi-sensor data fusion. The sensor models determine detection-based and prediction-based measurements via elliptical sampling (112) and take into account different uncertainties. In addition, the tracking approach exploits motion information by integrating displacements in the cost function for correspondence finding. Particles are detected by the spot-enhancing filter (SEF) (113) consisting of a Laplacian-of-Gaussian (LoG) filter followed by intensity thresholding of the filtered image and determination of local maxima.

### Motility analysis of ApoE^mT2^ and CD63^mCherry^

The motility of ApoE^mT2^- and CD63^mCherry^-positive puncta was quantified by a mean squared displacement (MSD) analysis (114) using the computed trajectories. For each trajectory with a minimum of 10 time points (corresponding to a time duration of 32.5 s), we computed the MSD as a function of the time interval *Δt*. All MSD curves corresponding to ApoE and CD63 respectively were averaged to obtain the respective MSD curves. To quantify the motility, we fitted the anomalous diffusion model *MSD(Δt) = 4ГΔt^α^* to the MSD values and obtained the anomalous diffusion exponent α for motion classification and the transport coefficient *Γ[µm^2^s^-α^].* The motion of ApoE^mT2^ and CD63^mCherry^ was classified into confined diffusion (α ≤ 0.1), obstructed diffusion (*0.1 < α < 0.9*), normal diffusion (*0.9 ≤ α < 1.1*), and directed motion (*α ≥ 1.1*) (115). To quantify the diffusion coefficient *D[µm^2^s^-1^],* we fitted the normal diffusion model *MSD(Δt) = 4DΔt* to the MSD values.

Automatic colocalization of ApoE^mT2^ and CD63^mCherry^ was performed using the computed trajectories with a minimum of 10 time points (corresponding to a time duration of 32.5 s). For each time point, colocalization was determined using a graph-based k-d-tree approach, which efficiently computes a nearest neighbor query based on Euclidean distances. An ApoE particle is considered to be colocalized with a CD63 particle, if the ApoE particle has a nearest CD63 particle within a maximum distance for at least a minimum number of consecutive frames. Otherwise, the ApoE particle is considered as non-colocalized with a CD63 particle. We used a maximum distance of 5 pixels (corresponding to 0.449 µm) and a minimum number of four consecutive frames (corresponding to 13 s). The computed colocalization information was visualized by color representations, and the motility of colocalized and non-colocalized ApoE was quantified by a MSD analysis.

To quantify the directed motion of colocalized ApoE^mT2^ and CD63^mCherry^, we performed a MSD analysis (114) using the computed colocalized trajectories of these proteins. To robustly classify the motion type into directed and non-directed motion of colocalized ApoE, we fitted for each trajectory the anomalous diffusion model *MSD(Δt) = 4ГΔt^α^* to the MSD values in two intervals from *Δt = 0* s to 25 s and from *Δt = 0* s to 60 s. Directed motion is considered if for one of the intervals we have α ≥ 1.1, otherwise non-directed motion is considered. For the classified trajectories, the MSD curves were averaged to obtain a MSD curve for colocalized ApoE with directed and non-directed motion, and the motion was quantified by the transport coefficient *Γ[µm^2^s^-α^]*, the diffusion coefficient *D[µm^2^s^-1^],* and the anomalous diffusion exponent α.

### Quantification and statistical analysis

Unless otherwise stated, differences between sample populations were evaluated using a two-tailed, unpaired Student’s *t-*test provided in the GraphPad Prism 8 software package. Differences with P-values less than 0.05 are considered to be significant and shown on the graph. The sample size of each experiment is specified in the corresponding figure legend.

## Acknowledgments

We thank Ulrike Herian, Stephanie Kallis, Marie Bartenschlager and Micha Fauth for excellent technical assistance and Fredy Huschmand for IT assistance. We thank Dr. Thomas Pietschmann at TWINCORE - Centre for Experimental and Clinical Infection Research (Hannover, Germany) for providing HEK293T-miR122 cells. We would like to acknowledge the microscopy support from the Infectious Diseases Imaging Platform (IDIP) at the Center for Integrative Infectious Disease Research (CIID, Heidelberg, Germany) and the University of Heidelberg Electron Microscopy Core Facility (EMCF, Heidelberg, Germany) headed by Dr. Stefan Hillmer. We thank Dr. Barbara Mueller, Djordje Salai and Thorsten Mueller (CIID, Heidelberg, Germany) for kindly providing the CLIPf, CLIP-ATTO590, and mScarlet constructs, respectively. Plasmids pTRE3G-NlucP and pCMV-Sport6-CD63-pHluorin were kind gifts from Masaharu Somiya (Addgene plasmid # 162595) and D.M. Pegtel (Addgene plasmid # 130901), respectively. We are grateful to all members of the Molecular Virology unit for continuous stimulating discussions.

## Supporting information captions

**S1 Fig. Functionality of ApoEmT2.**

(A) Validation of ApoE tagging with various fluorophores and confirmation of expression. Huh7-Lunet cells with stable knockdown (KD) of ApoE were transduced with lentiviruses encoding different fluorescently tagged-ApoE variants. After selection for stable expression, lysates of given cell pools were analyzed by Western blot using an ApoE-specific antibody. α-tubulin served as a loading control. mScarlet-C1: wildtype mScarlet; mScarlet-H: photo-stable mScarlet (M164H) variant. (B) Subcellular distribution of ApoE^mT2^ in HEK293T (left) and Hela cells (right) stably expressing this protein after lentiviral transduction and selection. Cells were characterized by confocal microscopy. (C-D) HCV replication in Huh7-Lunet/ApoE^mT2^ cells. (C) Cells were transduced with either an empty vector (Empty V), or wildtype ApoE (ApoE^wt^), or ApoE^mT2^, respectively, and selected for stable transgene expression. Cells were then electroporated with *in vitro* transcripts of the HCV Renilla luciferase (RLU)-reporter virus (JcR2a). HCV replication was determined at indicated time points by measuring RLU activities in cell lysates. (D) Amounts of core protein contained in cells from (C) at indicated time points were measured by chemiluminescence assay. Data are means from a representative experiment (n=3).

**S2 Fig. Functionality of ApoEmT2 as determined by rescue of infectious HCV particle production.**

(A-B) HCV replication in HEK293T-miR122-ApoE^mT2^ cells. (A) Cells were transduced with either an empty vector (Empty V), or wildtype ApoE (ApoE^wt^), or ApoE^mT2^, respectively, and electroporated with *in vitro* transcripts of the HCV Renilla luciferase (RLU)-reporter virus (JcR2a). HCV replication was determined at indicated time points by measuring RLU activities in cell lysates. RLU activities were normalized to the 4 h value to correct for the transfection efficiency. (B) Amounts of core protein contained in cells from (A) at indicated time points were measured by chemiluminescence assay. Data in both panels are means for a representative experiment (n=2). (C-D) Production of infectious HCV in HEK293T-miR122-ApoE^mT2^ cells. (C) At 24 and 48 h post-electroporation, amounts of extracellular core protein present in supernatants of cells from (A) were determined by chemiluminescence assay. (B) Culture supernatants harvested at 24 and 48 h post-electroporation were used to inoculate naïve Huh7.5 cells and HCV replication therein was measured by quantifying RLU activity at 72 h after inoculation. Virus titers normalized to HCV RNA replication in transfected cells are shown. Data in both panels are means for a representative experiment (n=2).

**S3 Fig. Colocalization of ApoE^mT2^ with Rab7 and ADRP.** Huh7-Lunet/ApoE^mT2^ cells were transduced with lentiviruses encoding Rab7^mCherry^ (upper panel) or ADRP^mCherry^ (lower panel). Cells were fixed and analyzed by confocal microscopy. Boxed areas in the left panels are shown as enlarged views in the panels on the right of each row. Arrowheads point to ApoE-Rab7 positive signals.

**S4 Fig. Characterization of ApoE variants and NS5A mutants.**

(A) ApoE-NS5A colocalization in cells replicating a full-length HCV genome. Huh7-Lunet/ApoE^mT2^ cells were electroporated with *in vitro* transcripts of the HCV genome Jc1. At 54 h p.e, cells were fixed, permeabilized, and incubated with NS5A- and PDI-specific antibodies for subsequent immunofluorescence staining. Images were acquired with a confocal microscope. Arrowheads: ApoE-NS5A signals. Note the high similarity to the structures detected in cells containing the split HCV genome (Figure 4). (B) An example of automatic detection of ApoE-NS5A double-positive puncta from (A). Circles and numbers mark the identity of each detected ApoE-NS5A structure. (C) Expression and secretion of SNAPf- and KDEL-tagged ApoE. Lysates and culture supernatants of Huh7-Lunet/ApoE-KD cells expressing ApoE^SNAPf^, or ApoE^mT2-KDEL^, or ApoE^KDEL^ were analyzed by Western blot with ApoE-specific antibody. β-actin served as a loading control. KDEL-tagged ApoE that is retained in the ER served as specificity control to determine ApoE^SNAPf^ secretion. (D) Expression of CLIPf-tagged NS5A. Huh7-Lunet cells were electroporated with RNA of the subgenomic replicon sgJFH1/NS5A^wt^ or sgJFH1/NS5A^CLIPf^, and cell lysates harvested at 24, 48, and 72 h p.e were analyzed by Western blot using NS5A-specific antibody. β-actin served as a loading control. (E) Colocalization of ApoE-NS5A double-positive structure with CD63. Huh7-Lunet/ApoE^SNAPf^ cells were electroporated with subgenomic replicon RNA encoding NS5A^CLIPf^ and after 72 h, cells were sequentially labeled with SNAP^SiR647^ and CLIP^ATTO590^ for 1 h, fixed, permeabilized, incubated with anti-CD63^AF488^ antibody, and subjected to confocal microscopy. Four images on the bottom show single or merged channels magnified views of the boxed area in the top overview image. Arrowheads point to ApoE-NS5A-CD63 triple-positive signals. (F) Secretion of NanoLuciferase (Nluc)-tagged NS5A. Huh7-Lunet cells were electroporated with RNA of the subgenomic replicon sgJFH1/NS5A^Nluc^, and cell lysates and supernatants harvested at 24, 48, and 72 h p.e were subjected to Nluc activity measurement. (G) Mitigation of ApoE-NS5A interaction by a mutation in NS5A domain I. HEK293T-miR122 cells were co-transfected with constructs encoding HA-tagged ApoE and either an empty vector, or myc-tagged NS5A^wt^, or myc-tagged NS5A^APK99AAA^, respectively. At 30 h p.t, cell lysates were subjected to immunoprecipitation (IP) using a myc-specific antibody and captured complexes were analyzed by Western blot with an HA-specific antibody. Band intensities of co-captured ApoE were quantified and values were normalized to the one obtained with NS5A^wt^ that was set to 1. Total cell lysate (0.5%) was loaded as input.

**S5 Fig. Detection of HCV RNA by single molecule (sm) FISH.**

(A) Schematic of the design of smFISH Hulu probes used to detect HCV RNA. These probes target a region encoding for NS3 (nucleotide 3733 - 4889 of the HCV JFH1 genome; GenBank accession number AB047639). (B) Specificity of HCV RNA detection by smFISH with Hulu probes. HCV RNA contained in Huh7-Lunet cells harboring a subgenomic replicon was detected by smFISH. Huh7-Lunet cells expressing the membrane sensor eYFP-CaaX (farnesylation signal from human HRAS protein) and that were used as recipient cells in co-culture experiments served as a negative control. (C) Detection of ApoE-associated HCV RNA in recipient cells. Huh7-Lunet/ApoE^mT2^/CD63^mCherry^ cells containing a subgenomic HCV replicon (donors) were co-cultured with Huh7-Lunet^eYFP-CaaX^ cells (recipients) for 24 h. Thereafter, cells were fixed and processed for visualization of HCV RNA by using smFISH. An overview image is shown on the top. Dashed area 1: donor cell; dashed area 2: recipient cell. Magnified views of indicated areas are shown on the bottom panels. Arrows point to ApoE-associated HCV RNA dots detected in both donor and recipient cells. Arrowheads indicate ApoE-negative HCV RNA dots in the recipient cell.

**S1 Table. Reagent or resource used in this study**

**S1 Movie. Intracellular co-trafficking of ApoE-CD63 complexes in an uninfected hepatocyte**

Huh7-Lunet/ApoE^mT2^ cells expressing CD63^mcherry^ were analyzed by live-cell time-lapse confocal microscopy. The trajectories of several ApoE-CD63 double-positive signals are marked. Frame interval = 3.61 sec. Duration of shown imaging = 111.91 sec.

S2 Movie. Secretion of an ApoE-associated CD63-positve intraluminal vesicle in an uninfected hepatocyte

Huh7-Lunet cells expressing ApoE^mT2^ and CD63^pHluorin^ were cultured in imaging medium (pH 7.4) and analyzed by live-cell time-lapse confocal microscopy with a focus on plasma membrane resident CD63-fluorescent signals. Frame interval = 3.14 sec. Duration of shown imaging = 200.96 sec.

S3 Movie. Uptake of donor-derived ApoE-CD63 complexes by a recipient cell

Donor (Huh7-Lunet/ApoE^mT2^/CD63^mCherry^) and recipient cells (Huh7-Lunet cells expressing eYFP-tagged CaaX) were co-cultured for 16 h and analyzed by live-cell time-lapse confocal microscopy. An area of a recipient cell (gray) showing the donor-derived ApoE-CD63 double-positive signals is shown. Frame interval = 3.81 sec. Duration of shown imaging: 118.11 sec.

S4 Movie. Long-term time-lapse confocal imaging of ApoE, NS5A, and E2 trafficking in a HCV-replicating hepatocyte

Huh7-Lunet cells stably expressing ApoE^mT2^ and C-NS2/E2^eYFP^ were electroporated with the replicon RNA encoding mCherry-tagged NS5A. Cells were subjected to time-lapse live-cell confocal microscopy to monitor ApoE, NS5A, and E2 signals from 5 to 54 h post-electroporation. A duration from 25.5 to 54 h is shown. Frame interval = 30 min.

S5 Movie. Abundance of ApoE-NS5A foci in a HCV-replicating hepatocyte

Huh7-Lunet cells stably expressing ApoE^mT2^ and C-NS2/E2^eYFP^ were electroporated with the replicon RNA encoding mCherry-tagged NS5A. At 48 h post-electroporation, cells were subjected to time-lapse live-cell confocal microscopy to monitor ApoE, NS5A, and E2 signals. Frame interval = 10.0 sec. Duration of shown imaging: 490.0 sec.

